# Dynamic evolution of AT-rich isochores shapes the genome of an intertidal fungus *Annulohypoxylon annulatoides*

**DOI:** 10.64898/2025.12.16.694779

**Authors:** Che-Yu Thiat-Ut Kuo, Yu-Ching Liu, Cheng-Ju Yang, Chieh-Ping Lin, Hung-Wei Chen, Huei-Mei Hsieh, Yu-Ming Ju, Tsui-Hua Liu, Yi-Chien Lee, Isheng Jason Tsai

**Affiliations:** Biodiversity Research Center, Academia Sinica, Taipei, Taiwan; Taipei Municipal Chien Kuo High School, Taipei, Taiwan; Institute of Plant and Microbial Biology, Academia Sinica, Taipei, Taiwan

## Abstract

Marine and coastal fungi experience intense environmental variability, yet the genomic mechanisms enabling filamentous fungi to tolerate such conditions remain unclear. From 56 fungal isolates collected along the Lailai rocky shore in northern Taiwan, we selected *Annulohypoxylon annulatoides* for deeper investigation due to its prevalence and distinctive stress response. Phenotypic assays revealed that this strain exhibits distinct growth and recovery dynamics under salinity, temperature, and UV stress compared to conspecific strains isolated from tree bark. To investigate the genomic basis of its adaptation, we generated a high-quality 41.8 Mbp *de novo* genome assembly with 11,529 predicted proteins. Across Hypoxylaceae genomes, we identified variably sized and dispersed AT-rich isochores, which in *A. annulatoides* were enriched in repeats and displayed low gene density. Despite differences in AT content, core gene content and Pfam domain profiles remained conserved. These AT-rich isochores exhibit several sequence and structural features consistent with scaffold/matrix attachment regions (S/MARs), raising the possibility that they influence higher-order genome organisation. Comparative analyses suggest they arose through independent repeat insertions or via ancestral repeat amplifications. Together, our findings point to a role for AT-rich isochores in shaping genome architecture and potentially mediating stress-responsive regulation, supporting the broader environmental flexibility observed in Hypoxylaceae, including adaptation to dynamic coastal habitats.

## Introduction

Marine and coastal fungi are increasingly recognised for their ecological roles and biotechnological potential (Bonugli-Santos et al., 2015). Life in the intertidal zone is shaped by sharp swings in solar radiation, salinity, and temperature, demanding tolerance to multiple concurrent stressors (El-Bibany et al., 2014; Pescheck et al., 2014). Among such habitats, rocky shores represent some of the most dynamic and challenging environments, characterised by fluctuating salinity, intermittent desiccation, and intense ultraviolet radiation (Johnson, 2024). Ongoing climate change is amplifying thermal, ultraviolet radiation, and osmotic extremes, posing additional challenges to the survival of many species (Bornman et al., 2014; Fan & McColl, 2024; Y. Huang et al., 2024; Vargas Zeppetello et al., 2022). Fungal tolerance to such conditions has therefore become a growing focus, particularly with respect to growth under increasingly harsh environments. Rocky shores, in particular, offer a tractable natural system for investigating species tolerance to the extreme environmental conditions expected under future climate change scenarios (Hawkins et al., 2020).

Despite these stressors, many fungi thrive in intertidal habitats. Early research has shown that numerous marine fungi are able to tolerate a wide range of temperatures and high salinity (Ritchie, 1957), while others develop thick, rigid, and carbonaceous fruiting bodies that likely buffer against desiccation and UV stress (González & Hanlin, 2010). Within this context, members of the family Hypoxylaceae (order Xylariales) stand out. Though typically known as forest endophytes or saprobes in tropical and subtropical regions (Franco et al., 2022; U’Ren et al., 2016), their discovery from Red Sea and mangrove ecosystems reveals their ecological plasticity (Abdel-Wahab, 2005). Genomic and transcriptomic studies further indicate that many Hypoxylaceae possess extensive secondary metabolite gene clusters (SMGCs), which may confer their tolerance to stressors (Franco et al., 2022; Overy et al., 2019). In addition, conserved stress-coping genes such as *Sln1, Hik1*, and *Sho1* have been identified in multiple Hypoxylaceae genomes (Abdel-Wahab, 2005), underscoring a genomic repertoire that may preadapt these fungi to survive the extreme variability of coastal environments.

We hypothesise that fungi inhabiting coastal environments possess genomic features and physiological traits shaped by repeated exposure to fluctuating and often severe stressors, including salinity shifts, thermal extremes, and ultraviolet radiation. To examine this hypothesis, we surveyed the Lailai rocky shore in northern Taiwan and identified *Annulohypoxylon annulatoides* (Hsieh et al., 2024), a member of the Hypoxylaceae. We characterised the phenotypic responses of this coastal isolate in comparison with five terrestrial (bark-derived) conspecific strains under varying temperature, salinity, and UV conditions, and generated a high-quality de novo genome assembly. We then placed these data in a comparative genomic framework spanning sixteen Hypoxylaceae species, with a focus on four *Annulohypoxylon* genomes. Importantly, while coastal habitats provide a strong ecological motivation for this study, this comparative approach allows us to distinguish environment-associated patterns from deeper lineage-wide features of genome evolution. Together, our analyses provide a framework for investigating how repeat-driven genome architecture evolves under environmental pressure and may contribute to ecological flexibility across fungal lineages.

## Results

### Hypoxylaceae dominate the fungal community of a rocky shore environment

We established four contiguous 25 × 25 m plots along the wave-cut platform of the Lailai rocky shore in Northern Taiwan (Areas 1 to 4; **Fig 1a**), spanning a gradient from the terrestrial vegetation line to the seaward edge. Within each area, three rock surfaces were randomly selected, and from each rock we sampled three distinct microhabitats –the upper surface, lateral sides, and rock crevices (**Fig. 1b**). A total of 56 fungal isolates were obtained and identified to at least the genus level. Members of Hypoxylaceae accounted for the majority of isolates (**Fig. 1c**), suggesting this family dominates the fungal assemblage at the Lailai rocky shore. Amongst the recovered taxa, *Annulohypoxylon* (three isolates), *Hypoxylon* (ten isolates)*, Penicillium* (nine isolates) and *Whalleya* (five isolates) were most frequently recovered.

**Figure 1.**
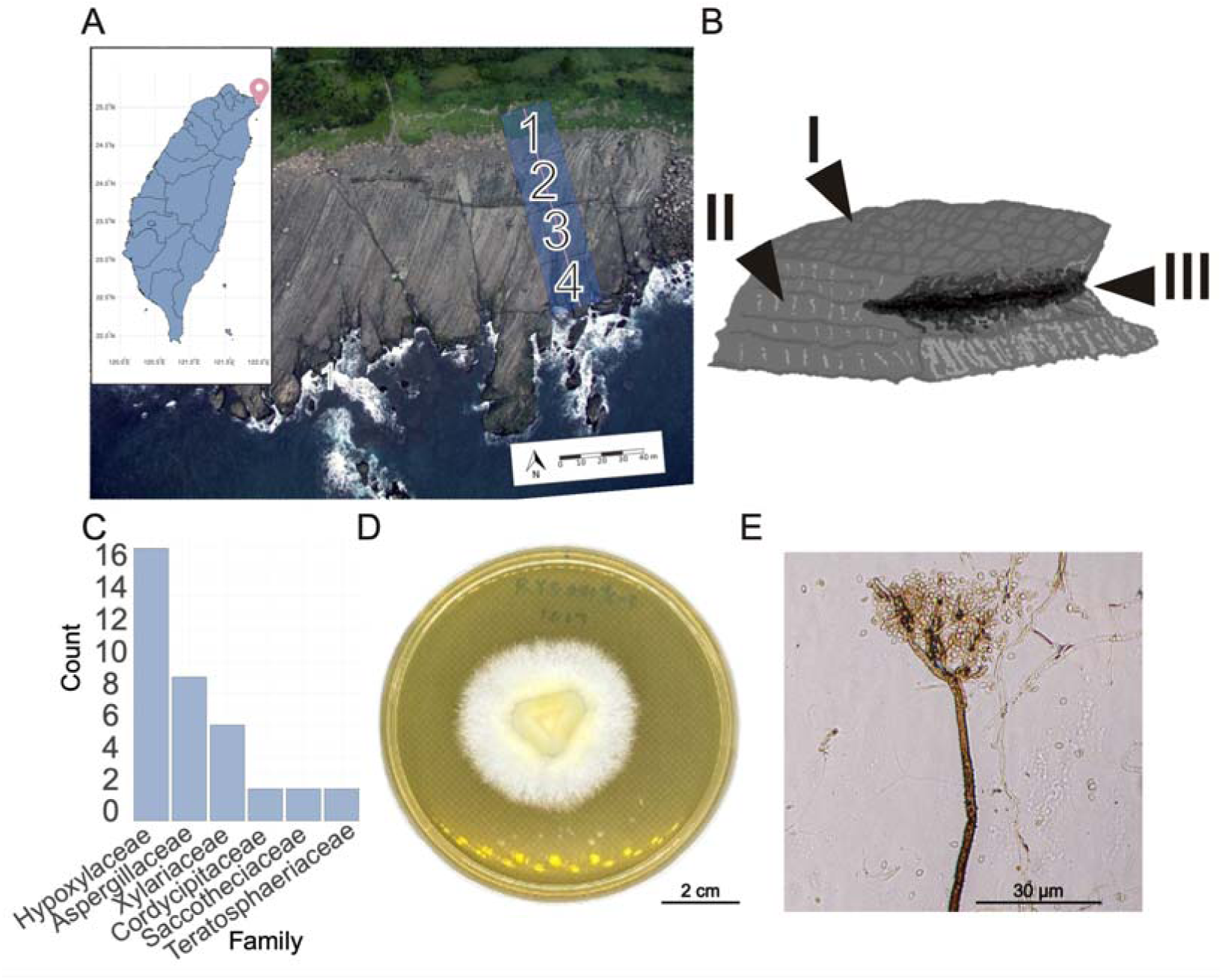
Sampling design, fungal diversity, and morphology of *A. annulatoides* RYS0019. (A) Aerial view of the Lailai rocky shore showing four sampling areas along a transect from the vegetation line (Area 1) to the coastline (Area 4) (photo: Cian-siang You, 2015). (B) Schematic of the three rock microhabitats sampled: (I) upper surface, (II) lateral side, and (III) crevice. (C) Number of isolates per fungal order, including only genera represented by more than two isolates. (D) Colony morphology and mycelial growth, and (E) conidiophore formation of *A. annulatoides* RYS0019 cultured on PDA at 30 °C.

Spatial comparisons showed Areas 1 and 4 yielded the highest number of isolates (16 each) yet exhibited distinct taxonomic profiles (**Supplementary Fig. 1a**), likely reflecting environmental gradients along the transect. Area 1 bordered terrestrial vegetation, whereas Area 4 was directly exposed to the shoreline. Among the three microhabitats examined, rock crevices supported the greatest fungal richness, harbouring the greatest number of isolates (22 isolates, **Supplementary Fig. 1b**). Hypoxylaceae species were consistently recovered across all areas and microhabitats, making Hypoxylaceae the dominant fungal group in this intertidal assemblage. Building on these observations, we investigated whether coastal Hypoxylaceae lineage exhibits genomic and phenotypic specialisation in response to coastal selective pressures. A representative strain was selected and identified as *Annulohypoxylon annulatoides* RYS0019 (**Fig. 1d**) based on internal transcribed spacer (ITS) phylogeny (**Supplementary Fig. 2**).

### Divergent stress recovery patterns in the coastal strain RYS0019 of *A. annulatoides*

We evaluated the stress tolerance of six *A. annulatoides* strains isolated from contrasting environment under varying temperature, salinity, and UV exposure conditions **(Supplementary Table S1)**. At a constant 30°C, bark-derived strains showed similar growth profiles (on average r = 5.40 mm²/hr and K= 549.00 mm²), while the marine strain RYS0019 exhibited similar average maximum growth rate (r = 5.69 mm²/hr) and carrying capacity (K = 408.96 mm^2^) (Fig. 2a). None of the strains grew at 400°C (**Supplementary Fig. 3 and 4**). Following UV exposure, most terrestrial strains recovered to near-control levels, whereas RYS0019 showed a pronounced decline in carrying capacity (*K* = 52.79 mm²) and maximum growth rate (r = 0.71 mm²/hr) (**Fig. 2b and Supplementary Fig. 5**). Under saline conditions, RYS0019 achieved optimal growth at 30‰, in contrast to other strains that showed reduced or unchanged growth at this concentration (**Supplementary Fig. 4 and 6**). These results indicate that strain RYS0019 exhibits a distinct stress-response profile compared to its forest counterparts, favouring resilience under moderate salinity but displaying slower growth and recovery under UV stress.

**Figure 2.**
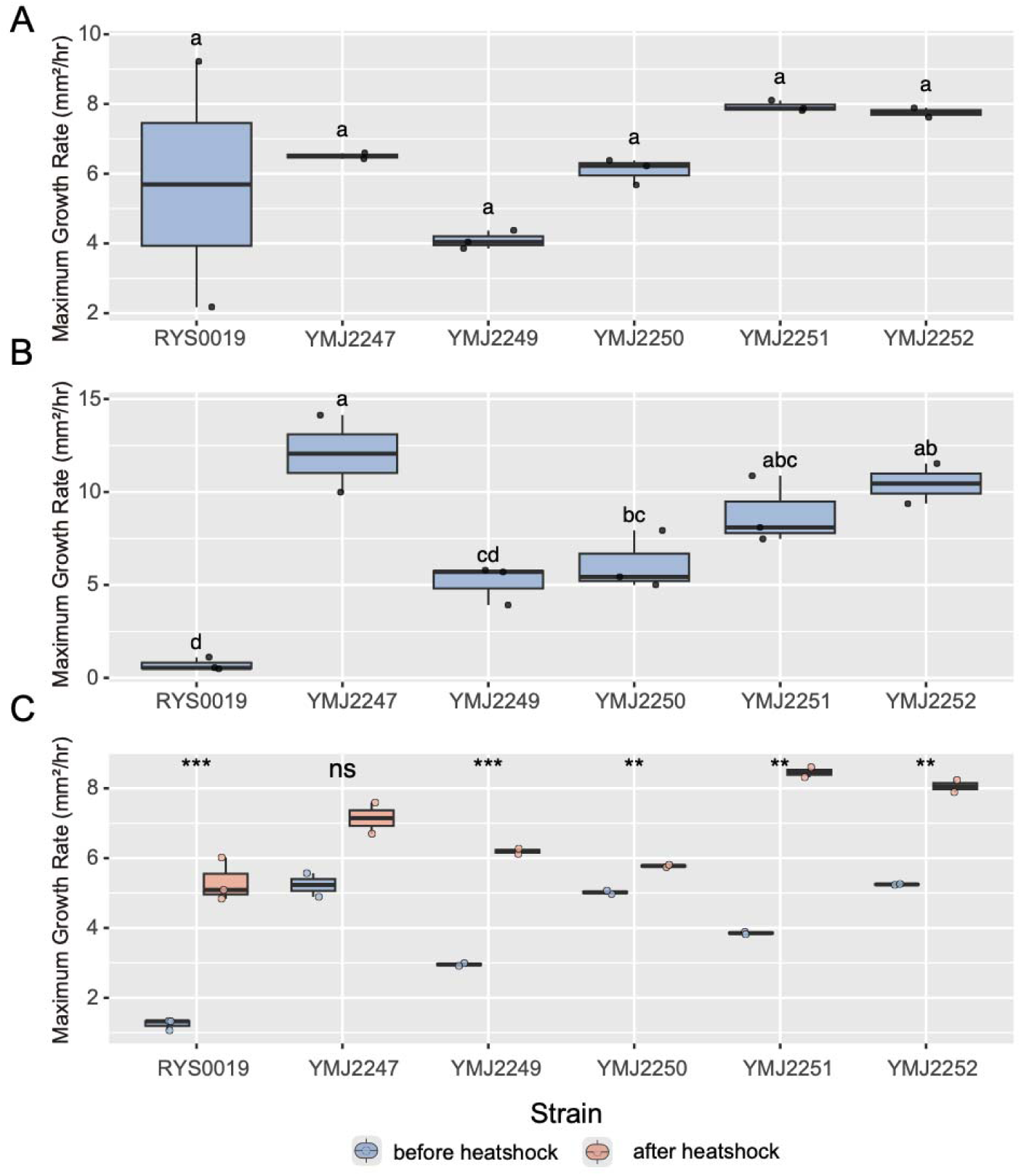
Growth performance of *A. annulatoides* strains under baseline and stress conditions. Maximum growth rates of six *A. annulatoides* strains measured under (A) standard growth at 30 °C in the dark, (B) after one day of UV shock followed by growth at 30 °C in the dark, and (C) before and after heat shock, in which cultures were exposed to 40 °C and then returned to 30 °C in the dark. Boxplots show median, interquartile range, and individual biological replicates. Different letters indicate significant differences among strains within each condition based on one-way ANOVA followed by Tukey’s post hoc test; asterisks denote significant differences between pre- and post-heat-shock treatments (ns, not significant; **P < 0.01; ***P < 0.001).

Building on its divergent stress responses, and observing that the ambient temperature at the time of sampling reached 32.3 °C, we next assessed whether RYS0019 exhibited a characteristic pattern of thermal recovery. Strains pre-grown under standard conditions were subjected to a one-day 40°C heat shock. All strains showed reduced growth during heat exposure, followed by increased growth rates post-treatment (**Fig. 2c**). In particular, RYS0019 displayed the most pronounced recovery, with its maximum growth rate increasing from 1.24 to 5.31 mm²/hr, potentially reflecting a strategy of rapid regrowth after transient stress. As all *A. annulatoides* strains are generally resilient to environmental fluctuation, these patterns suggest that RYS0019 prioritises post-stress recovery over maximal resistance, consistent with an environment characterised by frequent but transient stress episodes rather than sustained extremes.

### Genome assembly and annotation of *A. annulatoides* RYS0019

To investigate the genomic basis of environmental stress tolerance in *A. annulatoides*, we performed whole-genome sequencing of RYS0019 strain using Oxford Nanopore long-read technology. The initial assembly was produced with Flye (Kolmogorov et al., 2019) and subsequently polished with Illumina reads with NextPolish (Hu et al., 2020), resulting in a 41.8 Mbp genome assembly comprising 28 scaffolds with N50 of 4.4 Mbp (**Supplementary Table S2**). Gene prediction with the BRAKER3 (Gabriel et al., 2023) pipeline aided by reference protein homology support of closely related species and transcriptome sequencing identified 11,523 protein-coding gene models. Assembly completeness was estimated at 97.5% using Compleasm (N. Huang & Li, 2023), exceeding three other sequenced *Annulohypoxylon* species (**Supplementary Table S2**). Orthologue clustering with OrthoFinder (Emms & Kelly, 2019) on 16 Hypoxylaceae proteomes (**Supplementary Table S3**) and *Xylaria venustula* FL0490 as outgroup identified 12,686 orthologous groups (OGs), including 1,144 single-copy orthologues, of which 12,570 were shared across Hypoxylaceae species, while 119 were unique to Hypoxylaceae compared with outgroup *X. venustula* FL0490.

To determine the phylogenetic placement of *A. annulatoides,* we constructed a maximum-likelihood phylogeny based on a concatenated alignment of 1,144 single-copy orthologues and a coalescent-based phylogeny of individual gene trees (**Fig. 3a**). *A. annulatoides* RYS0019 clustered in a well-supported clade sister to *A. maeteangense* and *A. moriforme*. This lineage was resolved as sister to a second robust clade comprising *A. stygium*, *A. nitens*, *A.* sp. FPYF3050, and *Rhopalostroma terebratum*, together delineating two major groups within the *Annulohypoxylon* genus. The overall topology among *Hypoxylon* species was consistent with previous phylogenomic analyses of the order Xylariales (Franco et al., 2022), supporting the stability of these deep relationships within Hypoxylaceae.

**Figure 3.**
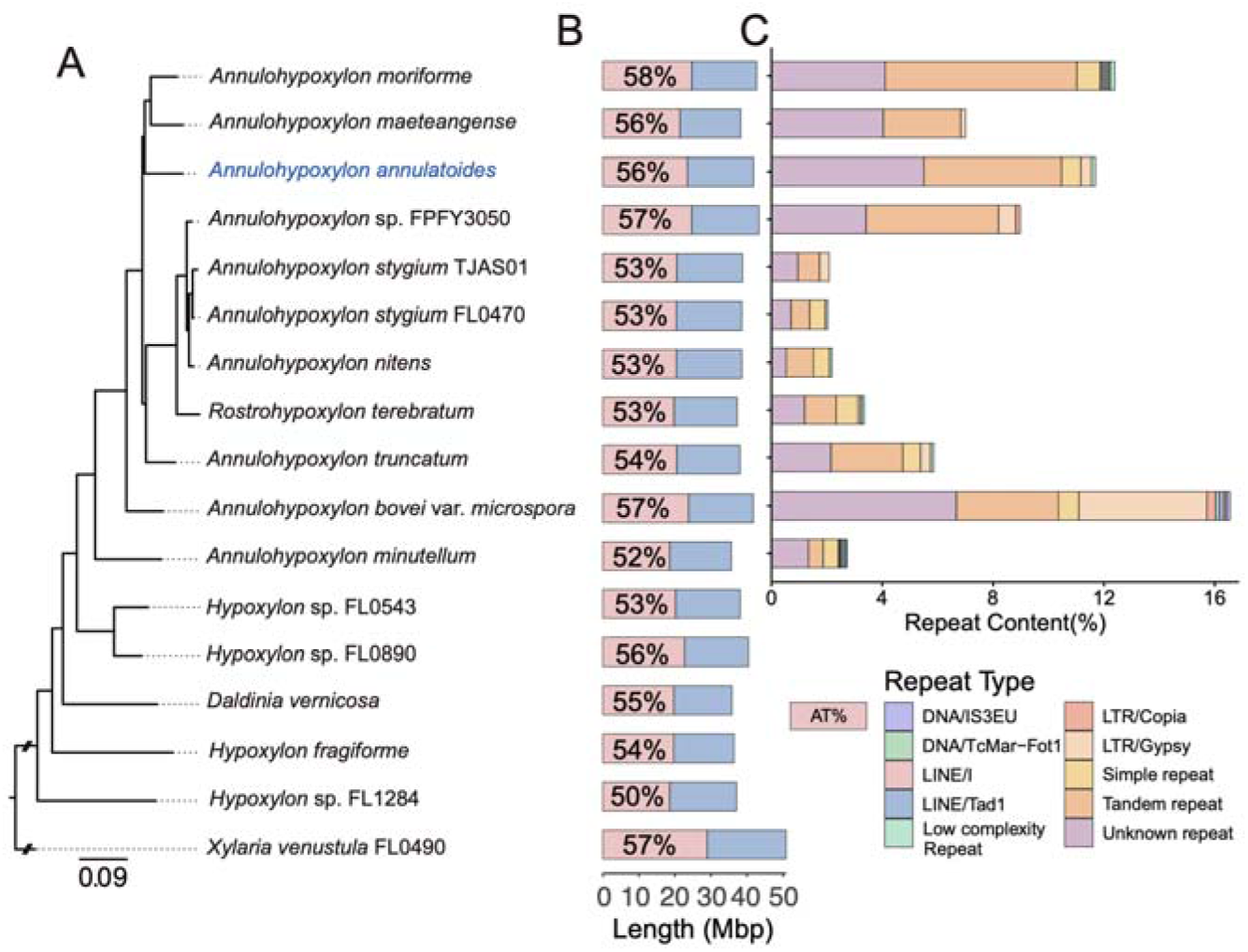
Phylogeny, genome size, and AT-rich repeat composition in Hypoxylaceae. (A) Phylogenetic relationships among 16 Hypoxylaceae species, rooted with *Xylaria venustula* FL0490 as the outgroup. (B) Genome size and overall AT content for each species. (C) Composition and relative abundance of major repeat classes across 11 *Annulohypoxylon* genomes.

Within the *Annulohypoxylon* clade, *A. annulatoides* possesses one of the largest genomes (41.8 Mbp; **Fig. 3b**), and AT content of 56%. *A. moriforme* and *A. maeteangense* formed a sister group with *A. annulatoides*, all of which possessed relatively large genomes ranging from 38.2 to 42.7 Mbp and high AT contents (56–58%) (**Fig. 3b**). The sister clade, comprising *A. stygium* TJAS01, *A. stygium* FL0470, and *A. nitens*, exhibited genome sizes between 38.5 and 38.8 Mbp, with lower AT contents of 53%. *A. truncatum* also belonged to this group, showing a genome size of 38.1 Mbp and an AT content of 54%. Outside these two lineages, *A. bovei* var. *microspora* displayed a genome size of 41.8 Mbp and an AT content of 57%, whereas *A. minutellum*, the most basal species in the phylogeny, had the smallest genome (35.7 Mbp) and the lowest AT content (52%).

### Repeat content and genome size

To account for the relatively larger genome size observed in *A. annulatoides*, we conducted repeat annotation across *Annulohypoxylon* genomes (**Fig. 3c**). Known repeats such as transposable elements (TEs) were not the predominant components. Instead, unknown and tandem repeats constituted the majority. In *A. annulatoides*, repetitive sequences spanned approximately 12% of the genome, with tandem and unknown repeats comprising approximately 11%, representing the highest proportions among all examined species. Similarly, several species with larger genomes including *A. maeteangense*, and *A.* sp. FPYF3050 also harboured elevated proportions of these repeat classes (6–11%).

Repeat content was positively correlated with genome size (Pearson’s r =0.78, P<0.001; **Supplementary Fig. 7**), indicating that repeat accumulation contributes substantially to genome expansion in this genus. AT content analyses revealed that repeats were consistently AT-rich across all species (**Supplementary Fig. 8**). Accordingly, genomes with greater repeat loads—including *A. annulatoides*, *A. maeteangense*, and *A.* sp. FPYF3050—displayed higher overall AT content (56–57%), whereas species with fewer repeats, such as *A. stygium* TJAS01, exhibited lower AT content (53%) (**Fig. 3c**). These patterns collectively indicate that AT-rich repetitive sequences are a major driver of genome expansion in *A. annulatoides* and related species.

### Repeats are mostly AT-rich isochores

Given the strong correlation between repeat abundance and genomic AT content, we next examined whether these repeats tend to accumulate within specific AT-rich compartments. Using OcculterCut segmentation (Testa et al., 2016), we identified extensive AT-rich tracts across all eight *Annulohypoxylon* genomes, ranging in size from 1,000 to 155,667 bp, with an average length of 11,102 bp—consistent with the hallmarks of AT-rich isochores (**Supplementary Fig. 9**). The majority of annotated repeats were localized within these regions: 69.9–86.8% of all repeats were located within AT-rich isochores, including 21.2–52.0% tandem repeats and 21.9–47.5% unknown repeats (**Supplementary Fig. 10a**). In *A. annulatoides*, some of these isochores contained homopolymeric tracts exceeding 29 identical bases (**Supplementary Fig. 11**). In contrast to their dense repeat load, AT-rich isochores exhibited markedly lower gene coverage—approximately 14-fold lower than GC-rich regions (**Supplementary Fig. 10b**), indicating that AT-rich isochores are predominantly intergenic, gene-poor genome compartments.

### Genome conservation and independent isochore origins

To assess whether isochores coincide with larger-scale structural changes, we investigated genome synteny among four *Annulohypoxylon* species with high-quality assemblies (N50 > 1 Mbp). Syntenic blocks were defined as regions in which the order and orientation of orthologous genes between two species were preserved, whereas syntenic breaks correspond to intergenic regions separating conserved blocks of orthologous genes. Comparisons between *A. annulatoides* and *A. maeteangense*, *A.* sp. *FPYF3050*, and *A. stygium* TJAS01 revealed that 90.23–91.54% of their genomes fell within conserved syntenic blocks, while having 165-217 syntenic breaks. Most syntenic blocks were assigned to the same orthologous chromosomes, indicating that despite the presence of numerous rearrangements, the majority of changes represent intra-chromosomal events (**Fig. 5a**).

Only a small fraction of breakpoints overlapped isochores: 13.33% in *A. maeteangense*, 12.15% in *A.* sp. *FPYF3050*, and 10.14% in *A. stygium*. Among inversion-associated breakpoints, overlap remained limited: 15.15% in *A. maeteangense*, 10.00% in *A.* sp. *FPYF3050*, and 6.52% in *A. stygium*. In total, just 1.64–2.60% of isochores were located at synteny breakpoints, indicating that the vast majority are embedded within regions of conserved gene order rather than associated with structural rearrangements. This pattern is visually evident in Figure 4a-b, where isochores frequently appear in intergenic regions flanked by conserved syntenic blocks. In addition, breakpoints containing isochores were substantially longer—averaging 43.63 to 65.65 kb—than those lacking isochores (mean length 9.36 to 11.86 kb), indicating that the presence of AT-rich isochores is associated with local intergenic expansion. These findings suggest that while most isochores arise independently within syntenically stable regions, a subset may insert into structurally dynamic loci and contribute to regional expansion.

**Figure 4.**
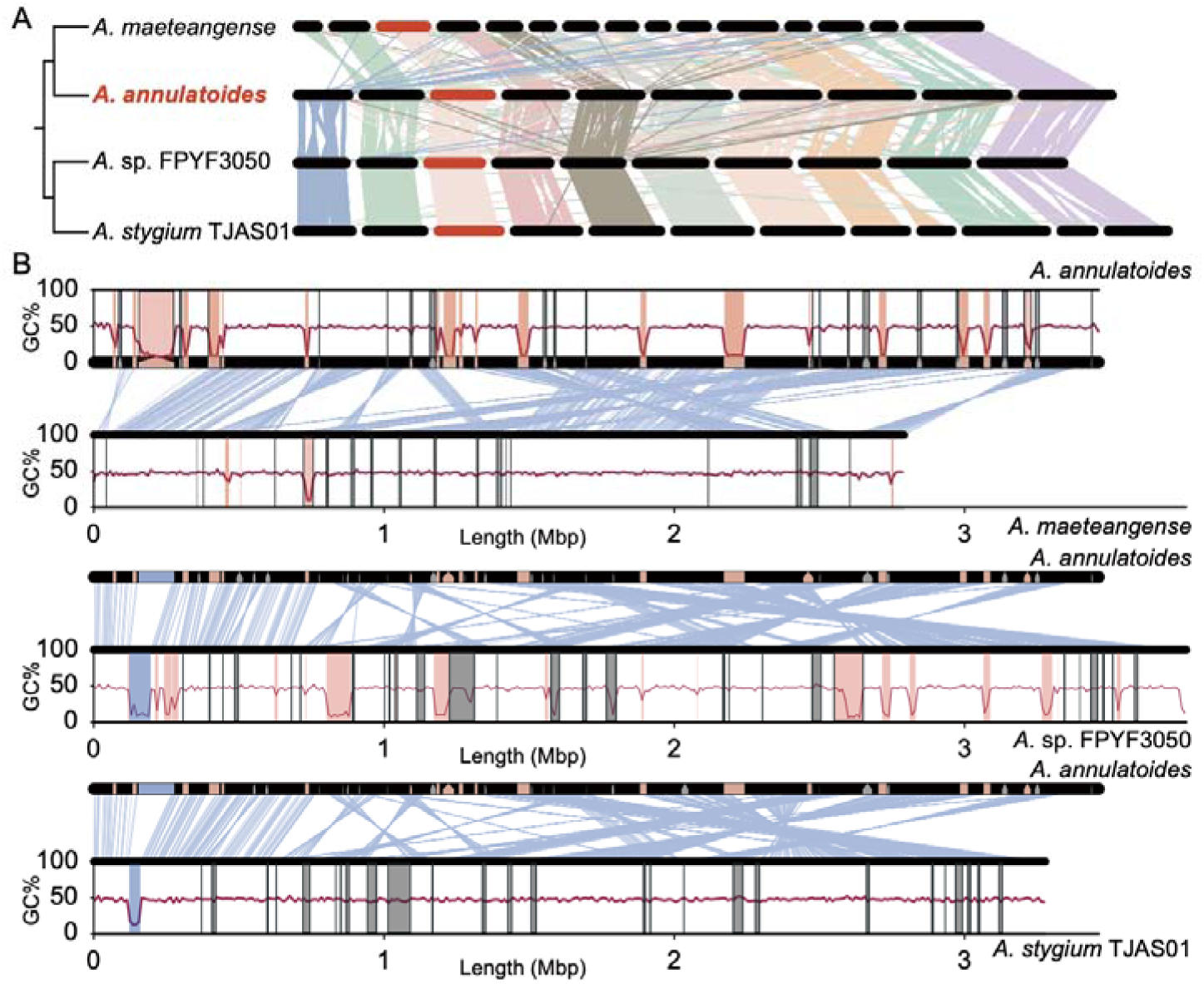
Synteny and intergenic isochore distribution among *Annulohypoxylon* species. (A) Syntenic relationships among chromosomes larger than 1 Mbp in *A. maeteangense*, *A. annulatoides*, *A.* sp. FPYF3050, and *A. stygium* TJAS01, defined using single-copy orthologs and shown as blue connecting lines. Scaffolds highlighted in red were extracted and shown in panel B for detailed comparison. (B) Distribution of intergenic regions containing isochores in *A. annulatoides* relative to the three comparison species. GC content is shown as a red line. Outlined rectangles or pentagons mark intergenic regions at syntenic breakpoints, with grey indicating absence and red indicating presence of isochores. In regions without block outlines, red denotes isochores unique to one species, whereas blue indicates isochores shared between corresponding intergenic regions. Genomic coordinates are shown in megabases (Mbp).

Across species, only 3.16%-10.27% of isochores were found in the same intergenic regions across species (**Supplementary Fig. 12**), indicating isochores are highly dynamic within otherwise conserved genome architecture. This is further illustrated in **Fig. 4b**, where conserved syntenic blocks are preserved across species, but the presence or absence of isochores at corresponding loci varies markedly. In *A. annulatoides*, *A. maeteangense*, *A.* sp. *FPYF3050*, and *A. stygium*, intergenic regions consisting of isochores were substantially longer than other orthologous intergenic regions (mean length 27.07–29.12 kbp vs. 1.95–2.08 kbp; **Fig. 5a**). Consistent with this, 76.09%, 87.50%, and 87.83% of intergenic regions that expanded by more than 10 kbp (relative to *A. maeteangense*, *A.* sp. FPYF3050, and *A. stygium* TJAS01) contained AT-rich isochores. In these cases, AT-rich isochores account for 82.23–85.71% of the total expansion, confirming that isochores are the primary drivers of large-scale intergenic extension.

**Figure 5.**
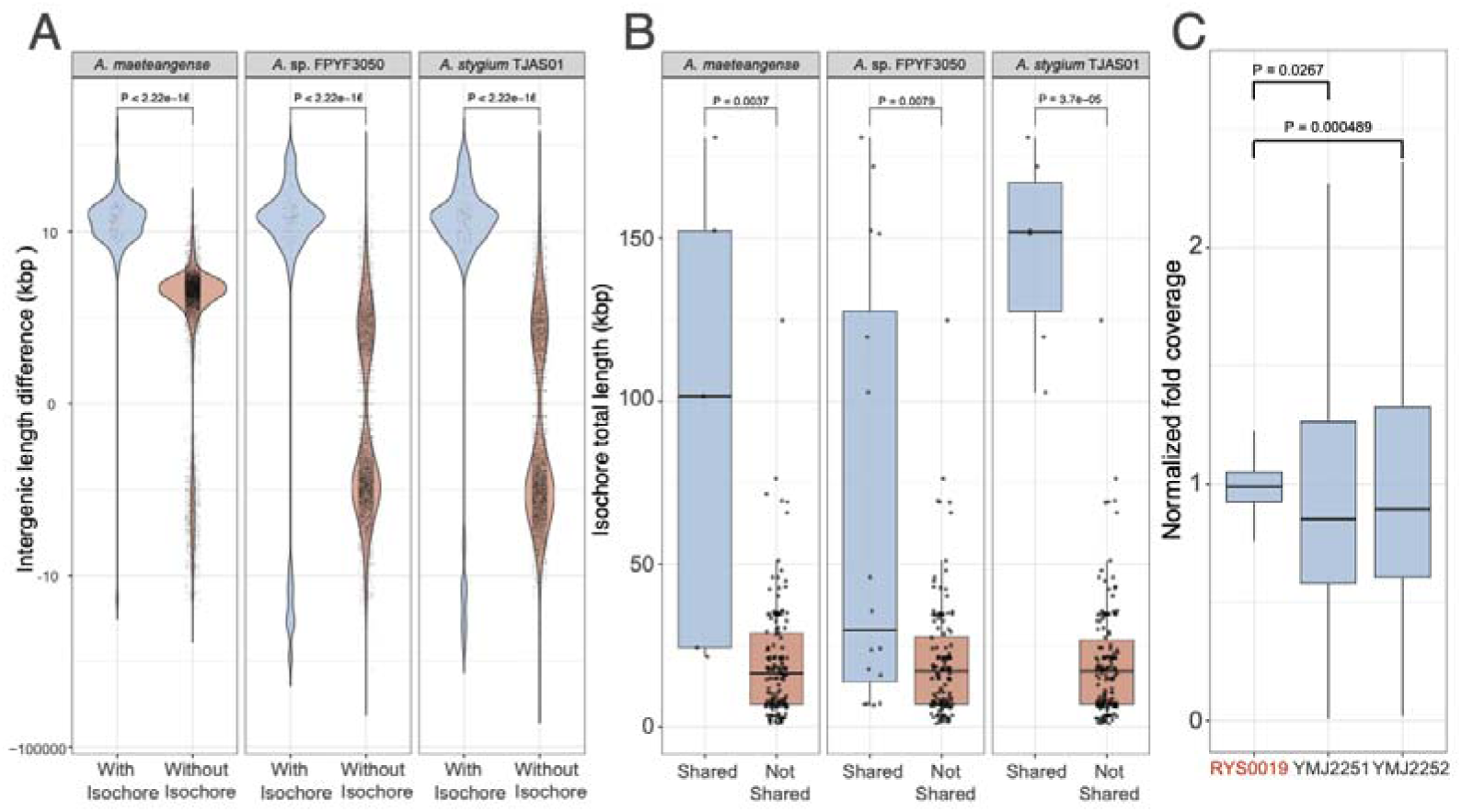
Intergenic expansion associated with AT-rich isochores in *Annulohypoxylon*. (A) Comparison of relative intergenic lengths of *A. annulatoides* compare with three related *Annulohypoxylon* species, contrasting intergenic regions containing AT-rich isochores with those lacking isochores. (B) Total lengths of AT-rich isochores in intergenic regions that are conserved between *A. annulatoides* and each comparison species (Shared) versus isochores located in intergenic regions where the corresponding species lacks an isochore (Not shared). (C) Normalized Illumina read coverage across AT-rich isochore regions in *A. annulatoides* RYS0019 and two terrestrial strains (YMJ2251 and YMJ2252). P values denote result from Wilcoxon Rank sum test.

A small subset of positionally conserved isochores was markedly longer than the remainder (**Fig. 4b**). For instance, in comparisons with *A. stygium*, shared isochores had a mean length of 14.64 kbp, whereas non-shared isochores averaged only 1.97 kbp (**Fig. 5b**). These unusually long, conserved intergenic intervals are consistent with centromeric or pericentromeric regions that are maintained across species, including lineages that otherwise lack extensive AT-rich isochores. Supporting this interpretation, resequencing analyses revealed pronounced differences in read coverage across isochore regions among *A. annulatoides* strains (**Fig. 5c**). The terrestrial strains YMJ2251 and YMJ2252 exhibited significantly reduced and more variable normalized coverage over AT-rich isochores relative to the coastal isolate RYS0019 (Wilcoxon rank-sum test, P < 0.05), indicating strain-specific divergence in isochore structure or copy number. Together, these results suggest that while a subset of long isochores is positionally constrained, the majority of AT-rich intergenic regions remain evolutionarily dynamic and differ substantially between marine and terrestrial lineages.

### Spatial and regulatory features of AT-rich isochores

To further investigate the distribution of AT-rich isochores, we examined their positions along chromosomes. Here, we compared the distance of each isochore from the chromosomal midpoint in *A. annulatoides* and *A.* sp. FPYF3050, the two species that exhibit strong AT-rich signals and share clear one-to-one chromosome correspondence. Kernel density estimates were generated to visualize the distribution of AT-rich isochores along chromosomes. In both species, isochores are enriched toward chromosome ends and are strongly depleted near the midpoint (**Supplementary Fig. 13a, b**).

To test whether isochore length varied by chromosomal position, we categorized isochores as near-terminal (distance to scaffold end ≤ 0.25) or central (> 0.25) and compared their lengths (**Supplementary Fig. 13c, d**). In *A. annulatoides*, median lengths between near-terminal and central isochores were nearly identical (15,989 vs. 13,871 bp). In *A.* sp. *FPYF3050*, the median was higher in central regions (21,290 bp) compared to terminal regions (11,752 bp), although the difference was not statistically significant (Wilcoxon test, p = 0.07). This result indicates that although AT-rich isochores consistently accumulate near chromosome ends across species, their lengths do not differ significantly across chromosomal regions. Despite variation in repeat composition and differences in length, the shared distribution patterns hint at conserved underlying processes.

To explore the functional potential of AT-rich isochores, we examined their spatial organisation and regulatory characteristics. These regions also showed a high density of predicted promoter motifs (**Supplementary Fig. 14**) across all six tested species and exhibited lower levels of DNA methylation than neighbouring GC-rich segments in *A. annulatoides* (**Supplementary Fig. 15**), indicating a more open chromatin configuration. However, genes located immediately downstream of AT-rich isochores were not significantly upregulated in *A. annulatoides* under normal culture conditions (**Supplementary Fig. 16**), suggesting that any regulatory influence of these regions is likely to be context-dependent rather than constitutive. Their low methylation and interspecies variability imply a latent regulatory capacity that may become active only under specific environmental or developmental contexts.

### Proteome specialisation of *A. annulatoides*

We compared the number of genes, protein families, transporters, and carbohydrate-active enzymes (CAZymes) across Hypoxylaceae species (**Supplementary Fig. 17b**). Protein families that were broadly enriched within Hypoxylaceae showed no further expansion in *A. annulatoides* RYS0019, suggesting that its adaptation does not involve further expansion of these gene families (**Supplementary Fig. 18**). Several stress-related protein families, however, were consistently enriched in Hypoxylaceae species compared to outgroup species. These included domains such as Ald_Xan_dh_C, associated with oxidative stress tolerance (Malik et al., 2019). Previously characterised stress-responsive genes, such as *Sln1*, *Hik1*, and *Sho1*, were also present in *A. annulatoides* but were not specifically expanded. These genes likely contribute to baseline resilience across both terrestrial and marine environments, though their expression dynamics under environmental stress remain to be investigated. Collectively, these findings suggest that *A. annulatoides* has retained the core genomic repertoire typical of Hypoxylaceae but may rely on regulatory or structural genome features—rather than gene content expansion—to achieve its environmental flexibility.

## Discussion

### Marine adaptation of *A. annulatoides*

Our survey revealed that Hypoxylaceae species dominated the fungal assemblage at the Lailai rocky shore, consistent with their broad ecological tolerance and previously reported presence in marine environments (Leman-Loubière et al., 2017; Schatz, 1988). Rock crevices harboured the highest number of Hypoxylaceae isolates (five isolates; **Supplementary Fig. 1b**), likely due to their relatively stable microclimatic conditions, including reduced desiccation and lower solar exposure. This widespread occurrence suggests that many members of the family possess an inherent capacity to persist under fluctuating conditions, likely supported by a conserved repertoire of stress-related genes. The coastal isolate *Annulohypoxylon annulatoides* RYS0019 exemplifies how such resilience has been further evolved to accommodate even greater environmental variability. Compared with its terrestrial counterparts, RYS0019 exhibited slower baseline growth but showed a much stronger recovery after transient heat stress. This pattern suggests a physiological adjustment that favors stability during environmental fluctuations—a strategy commonly observed among fungi in extreme environments (Gostinčar et al., 2022). At the genomic level, the identification of extensive AT-rich, the presence of extensive AT-rich, repeat-dense regions suggests that adaptation in A. annulatoides may extend beyond changes in gene content. Instead, variation in genome organisation itself may provide an additional layer of flexibility that complements conserved stress-response pathways.

### Evolutionary turnover of AT-rich isochores across species

Despite broadly conserved genome architecture across *Annulohypoxylon* species, AT-rich isochores exhibit striking evolutionary dynamism. The majority of isochores reside within syntenic regions, yet only a small fraction (1.64–2.60%) overlap synteny breakpoints, indicating that their emergence is largely decoupled from large-scale chromosomal rearrangements. The rare isochores that do coincide with structural breakpoints are more likely to have inserted after the formation of chromosomal rearrangements and subsequently expanded those breakpoint regions.

Isochores also show minimal positional conservation across species, with only 3.16–10.27% occupying homologous intergenic regions. This low degree of overlap suggests that most isochores were acquired independently in each lineage following species divergence. Nevertheless, a small subset of intergenic regions contains isochores that are conserved in both position and structure across species. These shared isochores are unusually long and may represent ancient elements inherited from a common ancestor. Their persistence, despite widespread lineage-specific gain and loss elsewhere in the genome, hints at potential functional constraints—possibly marking centromeric, pericentromeric, or structurally essential regions.

Taken together, these patterns support a model in which AT-rich isochores evolve rapidly and largely independently, driven by repeat accumulation within otherwise conserved genome regions. Their rare association with rearrangement sites and limited positional conservation across species argue against a scenario in which isochores are simply byproducts of structural genome evolution. Instead, the observations point toward post-divergence repeat-driven expansions as the primary mechanism driving isochore emergence and diversification.

### Possible mechanisms driving AT-rich isochore formation

Across *Annulohypoxylon* genomes, 70–87% of all annotated repeats are localized within AT-rich isochores, pointing to repeat accumulation as the primary driver of their formation. While repeat-rich regions in fungi are often attributed to transposable element (TE) activity (Zattera & Bruschi, 2022) or structural rearrangements (Xie et al., 2024), our synteny analyses show that isochores typically arise within conserved genomic contexts, independent of major chromosomal disruptions. Repeat annotation further revealed that most of these elements are tandem arrays or unknown repeats—not TE-derived—suggesting a distinct mutational origin.

Given their repeat composition, the initial formation of AT-rich isochores is likely seeded by slipped-strand mispairing, a mechanism known to promote tandem repeat expansion (Levinson & Gutman, 1987). However, the large size and widespread distribution of these isochores suggest that slippage alone is insufficient to explain their proliferation. Additional processes, such as unequal crossing-over (Smith, 1976) or replication-based amplification (Polleys et al., 2017), may have facilitated their expansion and dispersal across the genome. These mechanisms, acting recurrently and independently across lineages, could underlie the pervasive yet highly dynamic landscape of AT-rich isochores observed in *Annulohypoxylon* species.

### Putative Function of AT-rich isochores

The widespread presence of AT-rich isochores in *Annulohypoxylon* genomes suggests that genome architecture itself may play an active role in environmental adaptation. We propose two non-mutually exclusive mechanisms by which isochores may contribute to environmental adaptation.

First, the spatial distribution and sequence composition of isochores resemble scaffold/matrix attachment regions (S/MARs), which organize chromatin into loops by anchoring DNA to the nuclear matrix (Torres et al., 2023). S/MAR-like domains have been described in *Neurospora crassa* (Reckard et al., 2024), and if *Annulohypoxylon* isochores serve a similar role, their lineage-specific placement could reshape local chromatin topology (Torres et al., 2023). Such architectural restructuring may modulate the responsiveness of nearby genes—particularly those involved in stress tolerance—without requiring changes to the genes themselves. This type of topological flexibility could offer a mechanism for lineage-specific regulatory adaptation, especially under fluctuating environmental conditions (**Fig. 6**).

**Figure 6.**
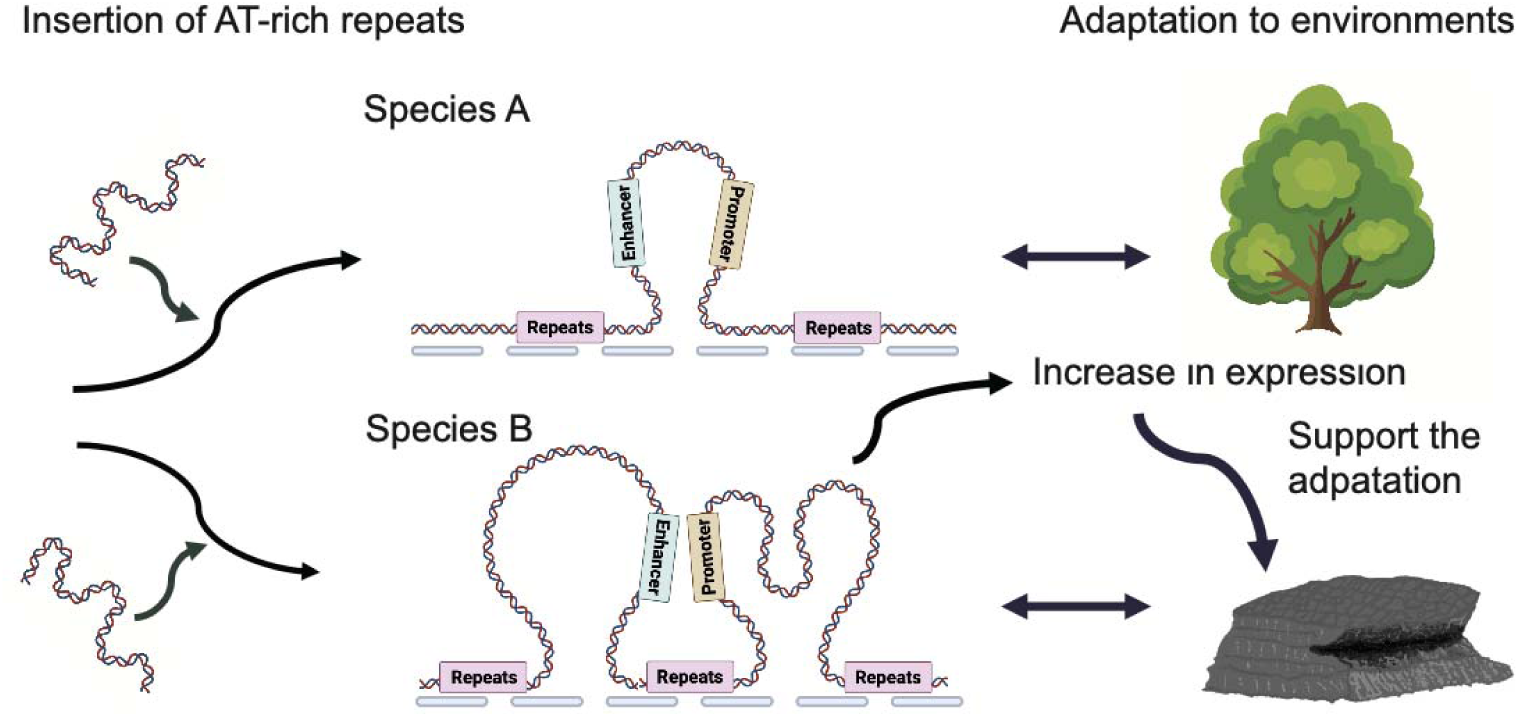
Hypothetical model illustrating how lineage-specific insertions of AT-rich sequences may contribute to environmental adaptation in Hypoxylaceae species. Species A: Promoter and enhancer elements are positioned within the same chromatin loop, limiting their interaction and preventing gene upregulation. Species B: Promoter and enhancer elements are compartmentalized into separate loops, facilitating interaction, leading to gene activation, and potentially enhancing stress tolerance.

Second, AT-rich isochores may act as latent cis-regulatory elements. Repetitive sequences can carry promoter- or enhancer-like motifs capable of influencing nearby transcription (Muszewska et al., 2019; Sawaya et al., 2013). In *Annulohypoxylon*, isochores are enriched in predicted promoter motifs and exhibit low levels of DNA methylation, suggesting an accessible chromatin state. Although adjacent genes were not broadly upregulated under standard laboratory conditions, these features suggest that isochores may function as inducible regulatory elements activated only under specific environmental or developmental contexts. This dynamic regulatory potential may complement conserved stress-response pathways, adding a flexible layer of transcriptional control.

The physiological profile of the coastal strain *A. annulatoides* RYS0019—marked by robust recovery following heat shock and preferential growth under moderate salinity—raises the possibility that isochores contribute to this resilience. This hypothesis, illustrated conceptually in Figure 6, provides a mechanistic link between genome structure and ecological plasticity and merits further investigation through chromatin conformation and gene expression studies under stress conditions.

## Conclusion

This study demonstrates that *Annulohypoxylon annulatoides* achieves environmental resilience through both physiological flexibility and genome architectural remodeling. The coastal isolate RYS0019 exhibits stress recovery traits suited to intertidal habitats, while comparative genomics reveals widespread, lineage-specific AT-rich isochores. These gene-poor, repeat-dense regions arise independently within conserved syntenic contexts and are largely uncoupled from chromosomal rearrangements. Their low methylation and structural resemblance to S/MARs suggest potential roles in shaping chromatin architecture and stress-responsive regulation. More broadly, this study highlights repeat-driven genome architecture as an underappreciated substrate for evolutionary innovation in fungi, offering a complementary route to ecological adaptation that operates independently of gene gain or loss.

## Materials and Methods

### Fungal isolation, culture, and strain identification

Fungal samples were collected from Lailai using sterile cotton swabs and immediately inoculated onto yeast extract–peptone–dextrose (YPD) agar supplemented with 30‰ salt. Plates were incubated at 20 °C until visible growth was observed. Individual colonies were then isolated, subcultured onto fresh YPD plates, and routinely maintained at 20 °C. Isolates that exhibited slow growth at 20 °C were subsequently transferred to 30 °C to facilitate optimal growth.

For strain identification, fresh mycelium from *A. annulatoides* cultures was scraped from the agar surface for genomic DNA and RNA isolation. Fungal DNA was extracted using both QuickExtract™ and the Quick-DNA™ Miniprep Kit, following manufacturers’ protocols. The internal transcribed spacer (ITS) region was amplified by PCR using ITS1F (5’-CTTGGTCATTTAGAGGAAGTAA-3’) and ITS4 (5’-TCCTCCGCTTATTGATATGC-3’) primers and Taq polymerase. PCR amplification was performed with an initial denaturation at 95 °C for 3 min, followed by 30 cycles of denaturation at 95 °C for 30 s, annealing at 52 °C for 30 s, and extension at 72 °C for 60 s, with a final extension at 72 °C for 5 min. When ITS amplification was unsuccessful, alternative sequence were amplified using primer pairs Bt2a (5’-GGTAACCAAATCGGTGCTGCTTTC-3’) and Bt2b (5’-ACCCTCAGTGTAGTGACCCTTGGC-3’) or CMD5 (5’-CCGAGTACAAGGAGGCCTTC-3’) and CMD6 (5’-CCGATAGAGGTCATAACGTGG-3’). PCR amplification was performed with an initial denaturation at 95 °C for 5 min, followed by 30 cycles of denaturation at 95 °C for 30 s, annealing at 50 °C (Bt2a/Bt2b) and 55 °C (CMD5/CMD6) for 30 s, and extension at 72 °C for 60 s, with a final extension at 72 °C for 5 min. PCR products were verified by agarose gel electrophoresis, subjected to Sanger sequencing, and the resulting sequences were compared against reference databases to confirm species identity.

### Strain phenotyping

Six *A. annulatoides* strains were analysed, comprising five terrestrial isolates (YMJ2247–YMJ2252) obtained from tree bark at distinct locations across Taiwan (Hsieh et al., 2024) and one coastal isolate, RYS0019, collected from rocky substrate at the Lailai rocky shore. To characterise phenotypic variation between marine and terrestrial strains, we employed two complementary approaches to quantify growth performance across a range of environmental conditions.

In the first approach, we examined the effects of temperature (20, 30, and 40 °C) and salinity (20‰, 30‰, and 70‰) by culturing strains on potato dextrose agar (PDA) and manually measuring colony diameters at two-day intervals. Each strain–condition combination was assayed with three biological replicates.

In the second approach, we captured fine-scale growth dynamics using an automated imaging and analysis pipeline. Colonies were scanned every 12 h using an Epson V850 scanner, and hyphal expansion was quantified with PyPhe (v0.98; Kamrad et al., 2020). Images were pre-processed in Photoshop, and hyphal area was segmented and quantified in FIJI using the Trainable Weka Segmentation plugin (Schindelin et al., 2012), trained and validated on 140 images each, achieving 98.33% classification accuracy. From these time-series data, maximum growth rates and carrying capacities were estimated in R.

The same automated workflow was applied to assess responses to acute environmental stress. For heat shock, cultures were grown at 30 °C for three days, exposed to 40 °C for one day, and then returned to 30 °C for a further six days. For UV shock, cultures were grown at 30 °C for three days, subjected to UV irradiation at 30 °C for one day, and subsequently transferred to PDA supplemented with 30‰ salt and incubated at 30 °C for six additional days. UV irradiation was conducted in a Class II Biological Safety Cabinet (11228 BBC 86) using a germicidal UV-C lamp (Sankyo Denki G30T8; peak wavelength 253.7 nm; UV radiant output 13.4 W). Sample was positioned 50 cm from the lamp surface. The UV dose was estimated to be ∼40.3 J/cm² over 24 hr. Colony growth throughout both shock treatments was monitored and quantified using the scanner-based pipeline described above. All assays were performed with three biological replicates per strain and condition.

### Library Preparation and Sequencing

Fresh mycelium from *A. annulatoides* cultures was collected from agar media for genomic DNA (gDNA) and RNA extraction. High-molecular-weight gDNA for Oxford Nanopore sequencing was isolated using a modified CTAB-based protocol, involving liquid nitrogen grinding, sequential chloroform extraction in the presence of isoascorbic acid, isopropanol precipitation, and RNase treatment as required, followed by DNA size selection and purification. Genomic DNA for Illumina (for *A. annulatoides* strain RYS0019, YMJ2251, and YMJ2252) sequencing was extracted using the Quick-DNA™ Microprep Kit.

The genome of *A. annulatoides* RYS0019 was sequenced using a combination of Oxford Nanopore long reads and Illumina short reads (BioProject PRJNA1381326). Oxford Nanopore sequencing was performed using a DNA library prepared with the ligation sequencing kit (SQK-LSK114; Oxford Nanopore Technologies, UK) and sequenced on an R10.4.1 flow cell (FLO-MIN114). Illumina short-read sequencing was outsourced for library preparation using the Illumina DNA PCR-Free Prep kit and sequenced on a NovaSeq X Plus platform. For gene model prediction and genome annotation, total RNA was extracted from *A. annulatoides* cultures using the Qiagen RNeasy Plant Mini Kit and sequenced on the Illumina platform.

### Genome Assembly

Oxford Nanopore reads were assembled de novo using Flye (v2.9.5; Kolmogorov et al., 2019) assembler. The resulting assemblies were first polished with Medaka (v1.11.3; https://github.com/nanoporetech/medaka) using Nanopore reads to improve consensus accuracy. Redundant haplotigs and overlapping contigs were subsequently identified and removed with Purge_Dups (v1.2.6; Guan et al., 2020). The curated assemblies were further refined by incorporating Illumina short reads using NextPolish (v1.4.1; Hu et al., 2020). Genome completeness was assessed with compleasm (v0.2.6; N. Huang & Li, 2023) using the ascomycota_obd10 fungi dataset.

### Phylogenetic Tree

Protein sequences from *Hypoxylon* sp. FL1284, *H. fragiforme*, *Hypoxylon* sp. FL0890, *Hypoxylon* sp. FL0543, *Daldinia vernicosa*, *Annulohypoxylon minutellum*, *A. bovei* var. *microspora*, *A. nitens*, *A. stygium* TJAS01, *A. stygium* FL0470, *Annulohypoxylon* sp. FPYF3050, *Rostrohypoxylon terebratum*, *A. truncatum*, *A. moriforme*, *A. maeteangense*, and *Xylaria venustula* FL0490 were retrieved from JGI and NCBI for phylogenetic reconstruction. Orthologous gene sets were identified using OrthoFinder (v2.5.5; Emms & Kelly, 2019). Concatenated amino acid alignments of single-copy orthologues were used to infer a maximum-likelihood phylogeny with IQ-TREE (v2.3.5; Minh et al., 2020).

### Repeat masking and Gene Annotation

In the genome assemblies of *A. stygium* TJAS01, *Annulohypoxylon* sp. FPYF3050, *A. maeteangense*, and *A. annulatoides*, tandem repeats were identified and soft-masked using Tandem Repeats Finder (v4.09.1; Benson, 1999). Additional classes of repetitive elements were detected and masked with EarlGrey (v4.2.4; Baril et al., 2024). Gene prediction was performed using BRAKER (v3.0.8; Gabriel et al., 2023), with the *A. annulatoides* hardmasked genome assembly, corresponding protein sequences, and RNA-seq data serving as inputs to train GeneMark-ETP (v1.0; Brůna et al., 2024) and AUGUSTUS (v3.5.0; Stanke et al., 2008). RNA-seq reads were mapped to the genome assembly using STAR (v2.7.11; Dobin et al., 2013), and the resulting alignments were employed to refine GeneMark. Protein files were retrieved from JGI Genome Portal (https://genome.jgi.doe.gov/portal/), included *A. bovei var. microspora*, *A. nitens*, *A. stygium*, *A. truncatum*, *A. moriforme*, and *A. maeteangense*, providing additional support for the prediction process.

Functional annotations of the amino acid sequences were carried out using eggNOG-mapper v2 (v2.1.2; default parameters; Huerta-Cepas et al., 2019) on eggNOG v5.0 database. Protein domains and families were identified using pfam_scan (v1.6; http://ftp.ebi.ac.uk/pub/databases/Pfam/Tools/) on Pfam database (release 37.2; Mistry et al., 2021). Protein sequence of *Hypoxylon* sp. FL1284, *H. fragiforme*, *H.* sp. FL0890, *H.* sp. FL0543, *Daldinia vernicosa*, *A. minutellum*, *A. bovei var. microspora*, *A. nitens*, *A. stygium* TJAS01, *A.* sp. FPYF3050, *Rostrohypoxylon terebratum*, *A. truncatum*, *A. moriforme*, *A. maeteangense*, *Xylaria venustula*, *Xylariaceae* sp. FL0821, *Xylaria digitata* 919, *Entoleuca mammata* CFL468, *Biscogniauxia nummularia*, *Fusarium avenaceum* MPI-SDFR-AT-0044, *Kretzschmaria zonata* GFP132, *Chaetomium tenue* MPI-SDFR-AT-0079, and *Nemania serpens* CBS 679.86 were used in comparison. We performed a Wilcoxon rank-sum test comparing domain counts between a defined Hypoxylaceae group (including *A. annulatoides*) and outgroup Ascomycota strain. Domains with Benjamini–Hochberg adjusted p-values < 0.05 were considered significantly enriched. The resulting domain abundance matrix was filtered to retain only enriched domains, and species labels were manually curated for downstream visualisation. Carbohydrate-active enzymes were annotated using run_dbcan (v2.0.11; https://github.com/linnabrown/run_dbcan).

### Genome analysis

Genome synteny was analysed among four *Annulohypoxylon* genomes based on orthologous gene relationships. Orthogroups were identified using OrthoFinder (v2.5.5; Emms & Kelly, 2019), and single-copy orthologues were extracted for pairwise species comparisons. Protein sequences from each species pair were compared using BLASTP (v2.15.0; Altschul et al., 1990), retaining only reciprocal single-copy orthologues. Syntenic blocks were subsequently inferred using DAGchainer (v1.12; (Haas et al., 2004), considering the linear order of orthologous genes along each genome while ignoring physical distance between loci. Promoters of *A. annulatoides*, *A. moriforme*, *A. maeteagense*, *A.* sp. *FPYF3050*, *A. truncatum*, and *A. bovei* var. *microspora* were performed using Promoter (v2.0; https://services.healthtech.dtu.dk/services/Promoter-2.0/).

AT-rich isochores were identified using OcculterCut segmentation (Testa et al., 2016). Downstream analyses focused on *Annulohypoxylon* species harbouring more than 1 Mbp of AT-rich sequence, as genomes with lower totals are more likely to reflect centromeric regions or insufficient assembly contiguity for robust inference. This criterion included all members of the clade comprising *A. annulatoides*, *A. moriforme*, and *A. maeteangense*, as well as *Annulohypoxylon* sp. FPYF3050 and *A. truncatum* from the sister clade, with *A. bovei* var. *microspora* included as an outgroup.

Genome-wide DNA methylation patterns in *A. annulatoides* were characterised using Oxford Nanopore reads. Raw FASTA files were processed with Dorado (v0.8.3; https://github.com/nanoporetech/dorado) and Modkit (v0.4.1; https://github.com/nanoporetech/modkit) under default parameters to detect 5-methylcytosine (5mC) and N6-methyldeoxyadenosine (6mA). Methylation calls supported by fewer than 10 reads or with posterior probabilities below 10% were excluded. Methylation levels were summarised in non-overlapping 10 kbp windows across the genome.

## Acknowledgement

We thank Chen Hsiao for advice on collection planning and Yu-Hsi Chuang for drawing the schematic diagrams in Figure 6.

## Author contributions

IJT, YiCL and CTK conceived the study. CTK, YuCL, CPL, and YiCL collected fungal samples. CTK and CJY extracted and sequenced the fungi genome. HMH and YMJ provided fungal strains *A. annulatoides* (isolates YMJ2247, YMJ2249–YMJ2252). CTK, HWC, YuCL, and YiCL analysed the genomics data. THL contributed to scientific discussions and provided intellectual input. CTK, YiCL, and IJT wrote the manuscript with input from all authors. All authors reviewed and approved the final version of the manuscript.

## Data Availability

All sequencing data generated in this study have been deposited in the NCBI Sequence Read Archive (SRA) under BioProject PRJNA1381326 and detailed in **Supplementary Table S1**.

## References

Abdel-Wahab, M. A. (2005). Diversity of marine fungi from Egyptian Red Sea mangroves. Botanica Marina, 48(5–6). 10.1515/bot.2005.047

Baril, T., Galbraith, J., & Hayward, A. (2024). Earl Grey: A Fully Automated User-Friendly Transposable Element Annotation and Analysis Pipeline. Molecular Biology and Evolution, 41(4). 10.1093/molbev/msae068

Benson, G. (1999). Tandem repeats finder: a program to analyze DNA sequences. Nucleic Acids Research, 27(2), 573–580. 10.1093/nar/27.2.573

Bonugli-Santos, R. C., dos Santos Vasconcelos, M. R., Passarini, M. R. Z., Vieira, G. A. L., Lopes, V. C. P., Mainardi, P. H., dos Santos, J. A., de Azevedo Duarte, L., Otero, I. V. R., da Silva Yoshida, A. M., Feitosa, V. A., Pessoa, A., & Sette, L. D. (2015). Marine-derived fungi: diversity of enzymes and biotechnological applications. Frontiers in Microbiology, 6. 10.3389/fmicb.2015.00269

Bornman, J. F., Barnes, P. W., Robinson, S. A., Ballaré, C. L., Flint, S. D., & Caldwell, M. M. (2014). Solar ultraviolet radiation and ozone depletion-driven climate change: effects on terrestrial ecosystems. Photochemical & Photobiological Sciences, 14(1), 88–107. 10.1039/c4pp90034k

Cian-siang You. (2015). Structural evolution study of lamprophyric dikes and country rocks in Lailai, northeastern coast of Taiwan. National Central University.

Dobin, A., Davis, C. A., Schlesinger, F., Drenkow, J., Zaleski, C., Jha, S., Batut, P., Chaisson, M., & Gingeras, T. R. (2013). STAR: ultrafast universal RNA-seq aligner. Bioinformatics, 29(1), 15–21. 10.1093/bioinformatics/bts635

El-Bibany, A. H., Bodnar, A. G., & Reinardy, H. C. (2014). Comparative DNA Damage and Repair in Echinoderm Coelomocytes Exposed to Genotoxicants. PLoS ONE, 9(9), e107815. 10.1371/journal.pone.0107815

Emms, D. M., & Kelly, S. (2019). OrthoFinder: phylogenetic orthology inference for comparative genomics. Genome Biology, 20(1), 238. 10.1186/s13059-019-1832-y

Fan, Y., & McColl, K. A. (2024). Widespread outdoor exposure to uncompensable heat stress with warming. Communications Earth & Environment, 5(1), 762. 10.1038/s43247-024-01930-6

Franco, M. E. E., Wisecaver, J. H., Arnold, A. E., Ju, Y., Slot, J. C., Ahrendt, S., Moore, L. P., Eastman, K. E., Scott, K., Konkel, Z., Mondo, S. J., Kuo, A., Hayes, R. D., Haridas, S., Andreopoulos, B., Riley, R., LaButti, K., Pangilinan, J., Lipzen, A., … U’Ren, J. M. (2022). Ecological generalism drives hyperdiversity of secondary metabolite gene clusters in xylarialean endophytes. New Phytologist, 233(3), 1317–1330. 10.1111/nph.17873

Gabriel, L., Brůna, T., Hoff, K. J., Ebel, M., Lomsadze, A., Borodovsky, M., & Stanke, M. (2023). BRAKER3: Fully automated genome annotation using RNA-seq and protein evidence with GeneMark-ETP, AUGUSTUS and TSEBRA. 10.1101/2023.06.10.544449

González, M. C., & Hanlin, R. T. (2010). Potential use of marine arenicolous ascomycetes as bioindicators of ecosystem disturbance on sandy Cancun beaches: Corollospora maritima as a candidate species. Botanica Marina, 53(6). 10.1515/bot.2010.073

Gostinčar, C., Zalar, P., & Gunde-Cimerman, N. (2022). No need for speed: slow development of fungi in extreme environments. Fungal Biology Reviews, 39, 1–14. 10.1016/j.fbr.2021.11.002

Guan, D., McCarthy, S. A., Wood, J., Howe, K., Wang, Y., & Durbin, R. (2020). Identifying and removing haplotypic duplication in primary genome assemblies. Bioinformatics, 36(9), 2896–2898. 10.1093/bioinformatics/btaa025

Hawkins, S. J., Pack, K. E., Hyder, K., Benedetti-Cecchi, L., & Jenkins, S. R. (2020). Rocky shores as tractable test systems for experimental ecology. Journal of the Marine Biological Association of the United Kingdom, 100(7), 1017–1041. 10.1017/S0025315420001046

Hsieh, H.-M., Lin, C.-R., Huang, C.-Y., & Ju, Y.-M. (2024). Annulohypoxylon annulatoides newly described in Taiwan. In Fung. Sci (Vol. 39, Issue 1).

Hu, J., Fan, J., Sun, Z., & Liu, S. (2020). NextPolish: a fast and efficient genome polishing tool for long-read assembly. Bioinformatics, 36(7), 2253–2255. 10.1093/bioinformatics/btz891

Huang, N., & Li, H. (2023). compleasm: a faster and more accurate reimplementation of BUSCO. Bioinformatics, 39(10). 10.1093/bioinformatics/btad595

Huang, Y., Liu, C., Huo, X., Lai, X., Zhu, W., Hao, Y., Zheng, Z., & Guo, K. (2024). Enhanced resistance to heat and fungal infection in transgenic Trichoderma via over-expressing the HSP70 gene. AMB Express, 14(1), 34. 10.1186/s13568-024-01693-5

Huerta-Cepas, J., Szklarczyk, D., Heller, D., Hernández-Plaza, A., Forslund, S. K., Cook, H., Mende, D. R., Letunic, I., Rattei, T., Jensen, L. J., von Mering, C., & Bork, P. (2019). eggNOG 5.0: a hierarchical, functionally and phylogenetically annotated orthology resource based on 5090 organisms and 2502 viruses. Nucleic Acids Research, 47(D1), D309–D314. 10.1093/nar/gky1085

Johnson, M. E. (2024). Ecology of Intertidal Rocky Shores Related to Examples of Coastal Geology across Phanerozoic Time. Journal of Marine Science and Engineering, 12(8), 1399. 10.3390/jmse12081399

Kamrad, S., Rodríguez-López, M., Cotobal, C., Correia-Melo, C., Ralser, M., & Bähler, J. (2020). Pyphe, a python toolbox for assessing microbial growth and cell viability in high-throughput colony screens. ELife, 9. 10.7554/eLife.55160

Kolmogorov, M., Yuan, J., Lin, Y., & Pevzner, P. A. (2019). Assembly of long, error-prone reads using repeat graphs. Nature Biotechnology, 37(5), 540–546. 10.1038/s41587-019-0072-8

Leman-Loubière, C., Le Goff, G., Retailleau, P., Debitus, C., & Ouazzani, J. (2017). Sporothriolide-Related Compounds from the Fungus *Hypoxylon monticulosum* CLL-205 Isolated from a *Sphaerocladina* Sponge from the Tahiti Coast. Journal of Natural Products, 80(10), 2850–2854. 10.1021/acs.jnatprod.7b00714

Levinson, G., & Gutman, G. A. (1987). Slipped-strand mispairing: a major mechanism for DNA sequence evolution. Molecular Biology and Evolution. 10.1093/oxfordjournals.molbev.a040442

Malik, N., Dhiman, P., Sobarzo-Sanchez, E., & Khatkar, A. (2019). Flavonoids and Anthranquinones as Xanthine Oxidase and Monoamine Oxidase Inhibitors: A New Approach Towards Inflammation and Oxidative Stress. Current Topics in Medicinal Chemistry, 18(25), 2154–2164. 10.2174/1568026619666181120143050

Minh, B. Q., Schmidt, H. A., Chernomor, O., Schrempf, D., Woodhams, M. D., von Haeseler, A., & Lanfear, R. (2020). IQ-TREE 2: New Models and Efficient Methods for Phylogenetic Inference in the Genomic Era. Molecular Biology and Evolution, 37(5), 1530–1534. 10.1093/molbev/msaa015

Mistry, J., Chuguransky, S., Williams, L., Qureshi, M., Salazar, G. A., Sonnhammer, E. L. L., Tosatto, S. C. E., Paladin, L., Raj, S., Richardson, L. J., Finn, R. D., & Bateman, A. (2021). Pfam: The protein families database in 2021. Nucleic Acids Research, 49(D1), D412–D419. 10.1093/nar/gkaa913

Muszewska, A., Steczkiewicz, K., Stepniewska-Dziubinska, M., & Ginalski, K. (2019). Transposable elements contribute to fungal genes and impact fungal lifestyle. Scientific Reports, 9(1), 4307. 10.1038/s41598-019-40965-0

Overy, D. P., Rämä, T., Oosterhuis, R., Walker, A. K., & Pang, K.-L. (2019). The Neglected Marine Fungi, Sensu stricto, and Their Isolation for Natural Products’ Discovery. Marine Drugs, 17(1), 42. 10.3390/md17010042

Pescheck, F., Lohbeck, K. T., Roleda, M. Y., & Bilger, W. (2014). UVB-induced DNA and photosystem II damage in two intertidal green macroalgae: Distinct survival strategies in UV-screening and non-screening Chlorophyta. Journal of Photochemistry and Photobiology B: Biology, 132, 85–93. 10.1016/j.jphotobiol.2014.02.006

Reckard, A. T., Pandeya, A., Voris, J. M., Gonzalez Cruz, C. G., Oluwadare, O., & Klocko, A. D. (2024). A constitutive heterochromatic region shapes genome organization and impacts gene expression in Neurospora crassa. BMC Genomics, 25(1), 1215. 10.1186/s12864-024-11110-7

Ritchie, D. (1957). Salinity Optima for Marine Fungi Affected by Temperature. American Journal of Botany, 44(10), 870. 10.2307/2438907

Sawaya, S., Bagshaw, A., Buschiazzo, E., Kumar, P., Chowdhury, S., Black, M. A., & Gemmell, N. (2013). Microsatellite Tandem Repeats Are Abundant in Human Promoters and Are Associated with Regulatory Elements. PLoS ONE, 8(2), e54710. 10.1371/journal.pone.0054710

Schatz, S. (1988). Hypoxylon oceanicum sp.nov. from mangroves. Mycotaxon, 33, 413–418.

Schindelin, J., Arganda-Carreras, I., Frise, E., Kaynig, V., Longair, M., Pietzsch, T., Preibisch, S., Rueden, C., Saalfeld, S., Schmid, B., Tinevez, J.-Y., White, D. J., Hartenstein, V., Eliceiri, K., Tomancak, P., & Cardona, A. (2012). Fiji: an open-source platform for biological-image analysis. Nature Methods, 9(7), 676–682. 10.1038/nmeth.2019

Testa, A. C., Oliver, R. P., & Hane, J. K. (2016). OcculterCut: A Comprehensive Survey of AT-Rich Regions in Fungal Genomes. Genome Biology and Evolution, 8(6), 2044–2064. 10.1093/gbe/evw121

U’Ren, J. M., Miadlikowska, J., Zimmerman, N. B., Lutzoni, F., Stajich, J. E., & Arnold, A. E. (2016). Contributions of North American endophytes to the phylogeny, ecology, and taxonomy of Xylariaceae (Sordariomycetes, Ascomycota). Molecular Phylogenetics and Evolution, 98, 210–232. 10.1016/j.ympev.2016.02.010

van Wyk, S., Wingfield, B. D., De Vos, L., van der Merwe, N. A., & Steenkamp, E. T. (2021). Genome-Wide Analyses of Repeat-Induced Point Mutations in the Ascomycota. Frontiers in Microbiology, 11. 10.3389/fmicb.2020.622368

Vargas Zeppetello, L. R., Raftery, A. E., & Battisti, D. S. (2022). Probabilistic projections of increased heat stress driven by climate change. Communications Earth & Environment, 3(1), 183. 10.1038/s43247-022-00524-4

Xie, F.-Y., Zhang, X.-G., Chen, J., Xu, X., Li, S., Xia, T.-J., Chen, L.-N., Yin, S., Ou, X.-H., & Ma, J.-Y. (2024). Downstream transcription promotes human recurrent CNV associated AT-rich sequence mediated genome rearrangements in yeast. IScience, 27(12), 111508. 10.1016/j.isci.2024.111508

Zattera, M. L., & Bruschi, D. P. (2022). Transposable Elements as a Source of Novel Repetitive DNA in the Eukaryote Genome. Cells, 11(21), 3373. 10.3390/cells11213373

