## Supplementary Information for "Dynamic evolution of AT-rich isochores shapes the genome of an intertidal fungus *Annulohypoxylon annulatoides*"

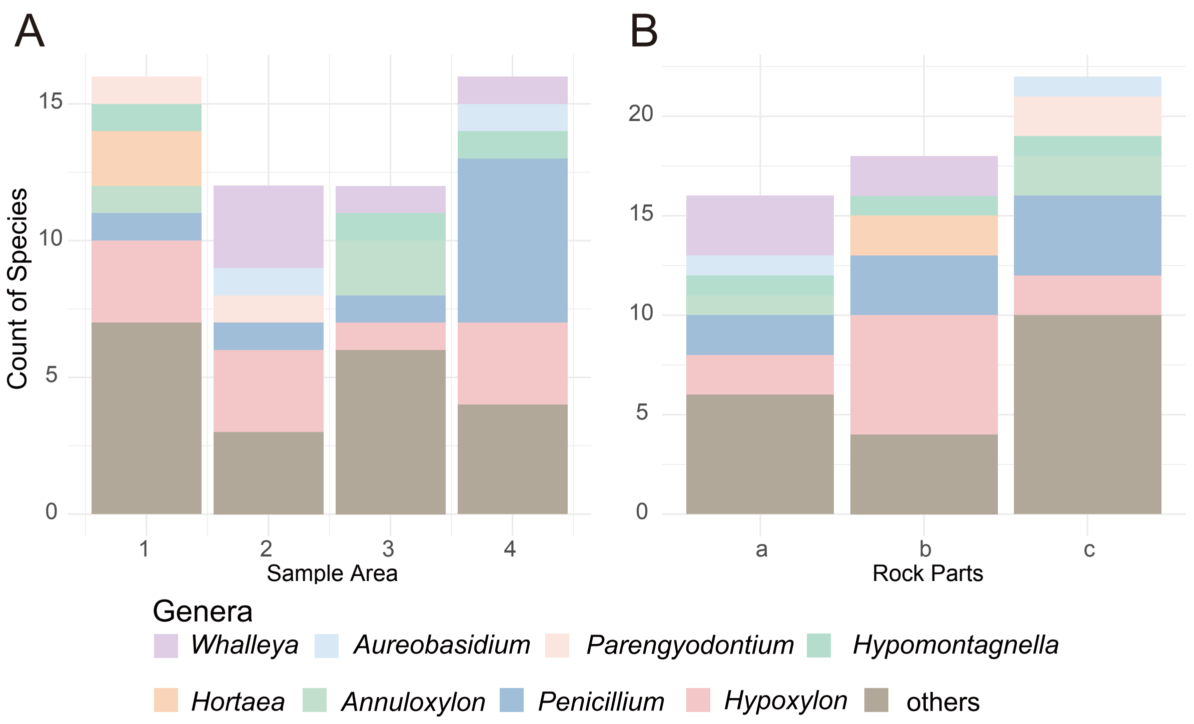


**Supplementary Figure 1. Genus-level composition of culturable fungi across the Lailai rocky shore sampling design.** (A) Stacked bar plots showing the number of isolates recovered per genus in each sampling area (Areas 1–4 along the transect from the vegetation line to the seaward edge). (B) Stacked bar plots showing the number of isolates recovered per genus across the three sampled rock microhabitats (a–c; corresponding to the upper surface, lateral side, and crevice, respectively). Each coloured segment denotes a genus, and bar height reflects total isolate counts per category.

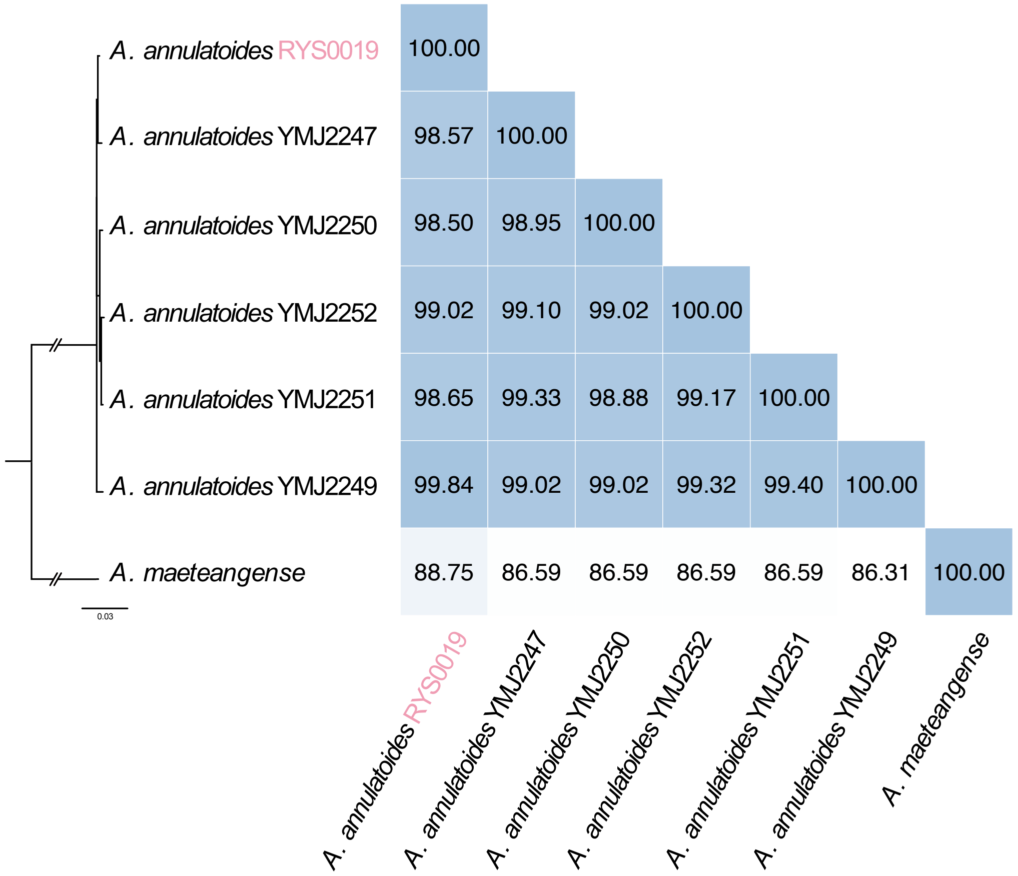


**Supplementary Figure 2. Phylogenetic relationships and pairwise sequence similarity among six *Annulohypoxylon annulatoides* strains.**

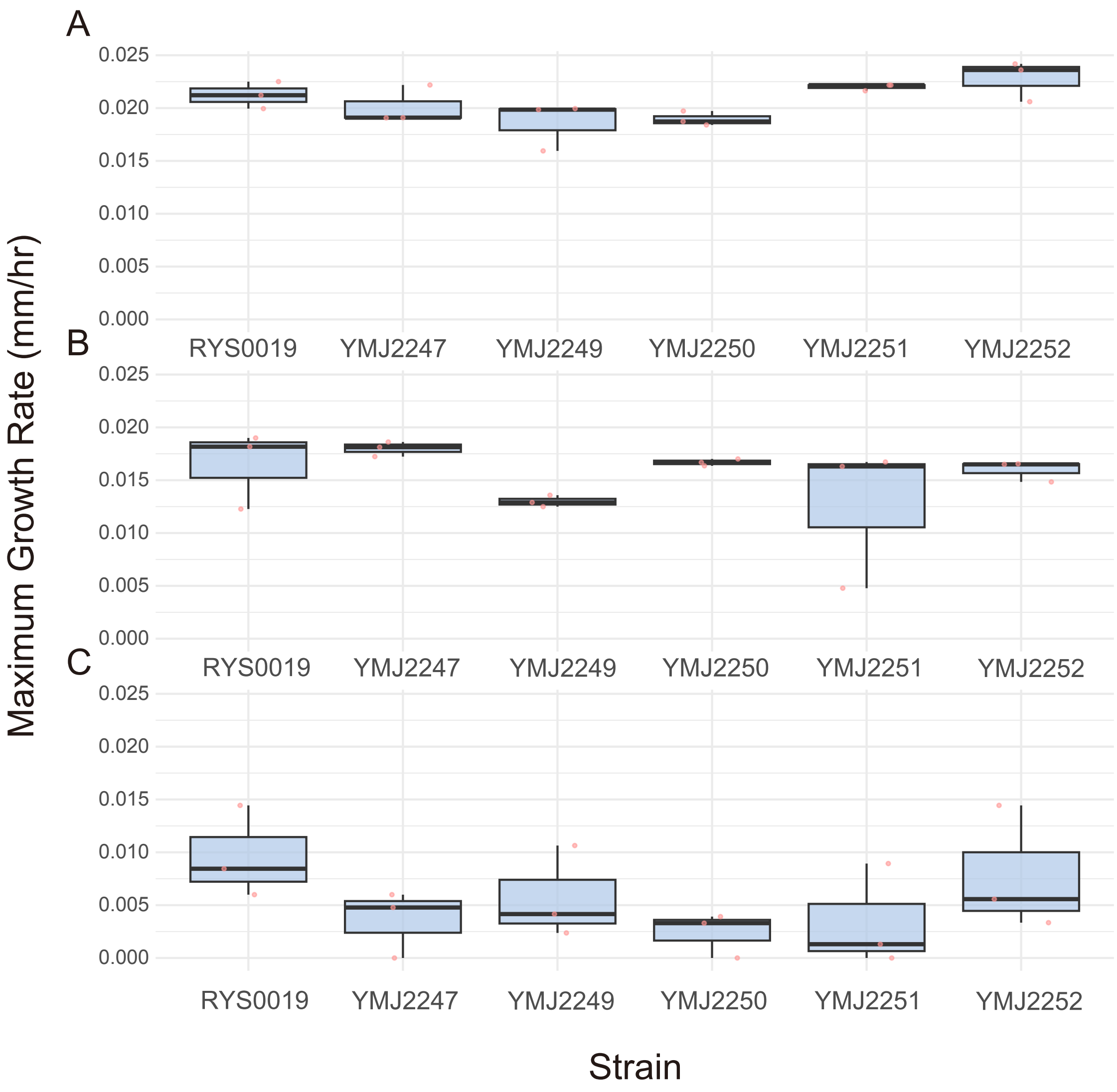


**Supplementary Figure 3. Temperature-dependent growth performance of six *Annulohypoxylon annulatoides* strains.** Boxplots show maximum growth rates of the coastal strain RYS0019 and five terrestrial strains (YMJ2247–YMJ2252) cultured in 0‰ NaCl medium at (A) 20 °C, (B) 30 °C, and (C) 40 °C. Points indicate individual biological replicates.

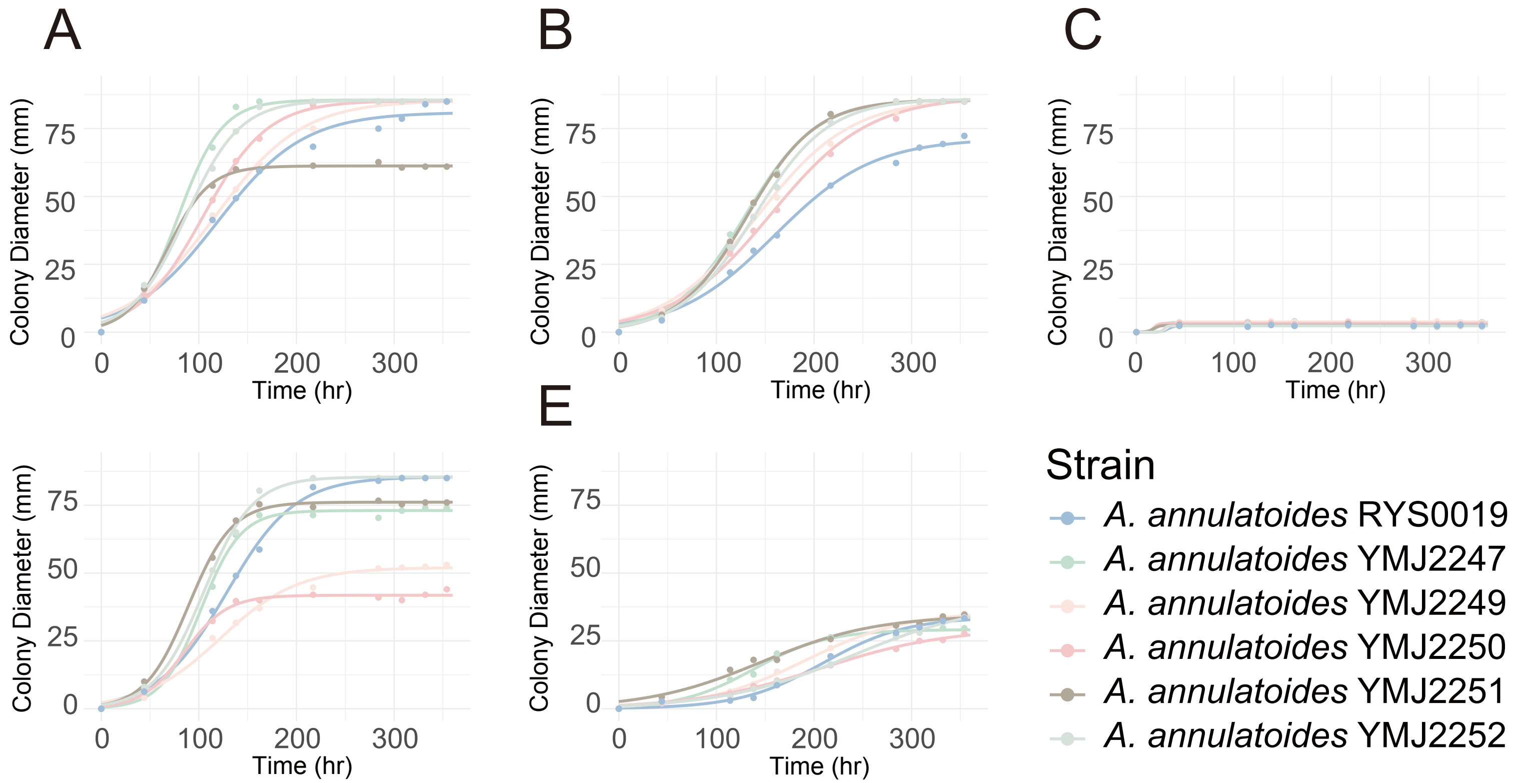


**Supplementary Figure 4. Growth dynamics of six *Annulohypoxylon annulatoides* strains under varying temperature and salinity conditions.** Time-course growth curves show hyphal expansion of the coastal strain RYS0019 and five terrestrial strains (YMJ2247–YMJ2252) cultured under five environmental conditions: (A) 30 °C, 0‰ NaCl; (B) 20 °C, 0‰ NaCl; (C) 40 °C, 0‰ NaCl; (D) 30 °C, 30‰ NaCl; and (E) 30 °C, 70‰ NaCl. Curves represent mean growth trajectories across biological replicates.

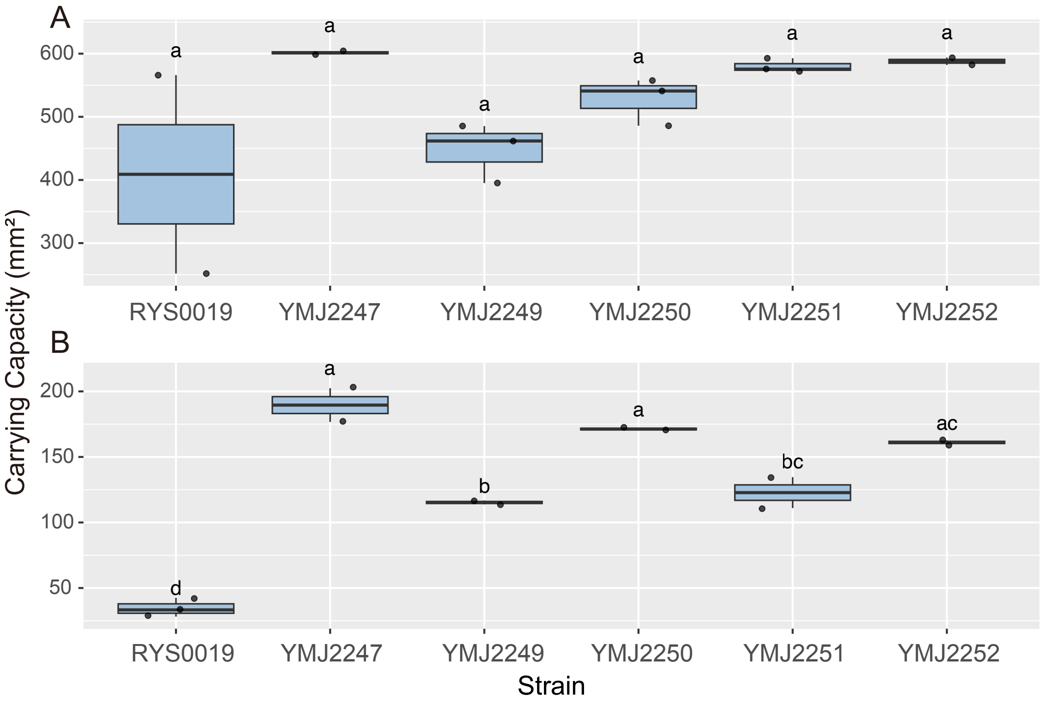


**Supplementary Figure 5. Effects of UV stress on carrying capacity in six *Annulohypoxylon annulatoides* strains.** Boxplots show carrying capacity (mm²) of the coastal strain RYS0019 and five terrestrial strains (YMJ2247–YMJ2252) under two conditions: (A) standard growth at 30 °C in 0‰ NaCl medium, and (B) following a one-day UV shock, with subsequent recovery at 30 °C in 0‰ NaCl medium. Letters above boxplots indicate statistically significant groupings based on one-way ANOVA followed by Tukey's post hoc test ($P < 0.05$).

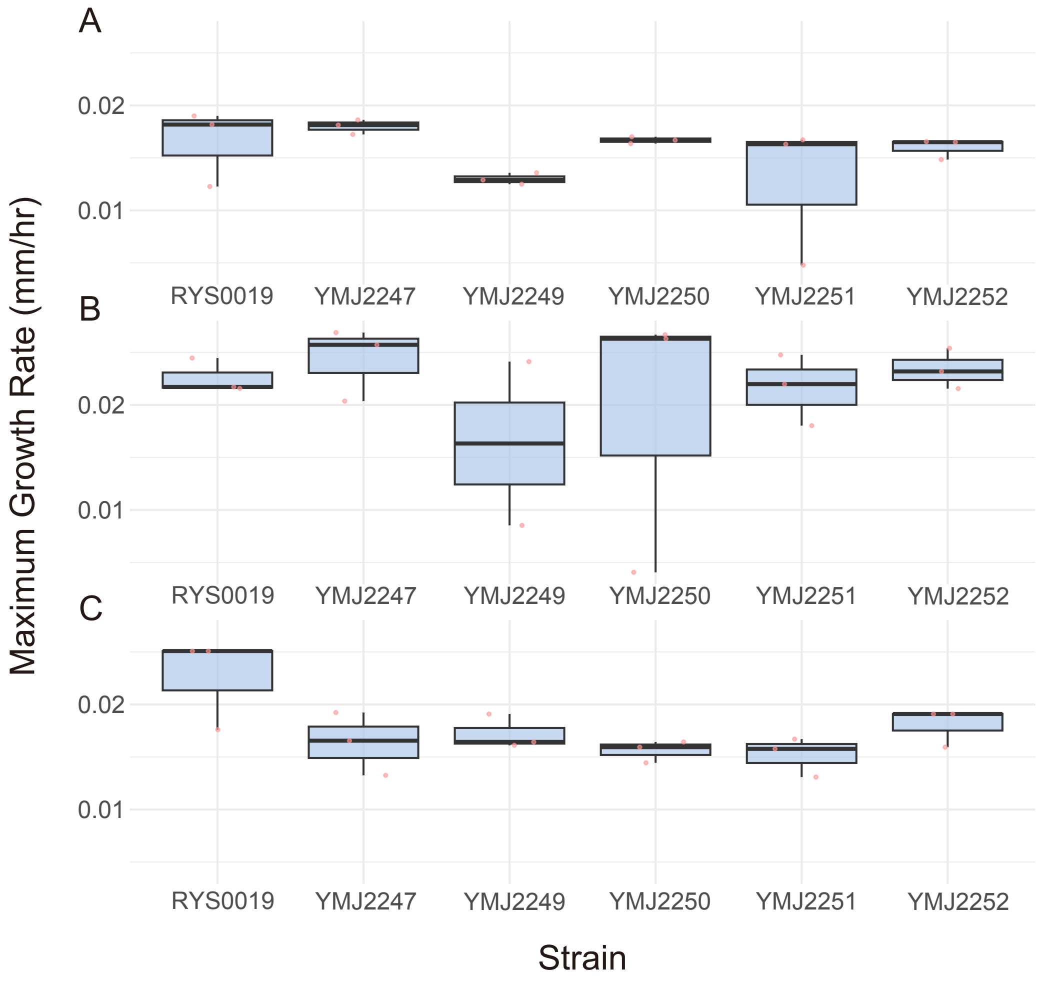


**Supplementary Figure 6. Salinity-dependent growth performance of six *Annulohypoxylon annulatoides* strains at 30 °C.** Boxplots show maximum growth rates ($mm^2\ hr^{-1}$) of the coastal strain RYS0019 and five terrestrial strains (YMJ2247–YMJ2252) cultured at 30 °C under three salinity conditions: (A) 0‰ NaCl, (B) 30‰ NaCl, and (C) 70‰ NaCl. Points represent individual biological replicates.

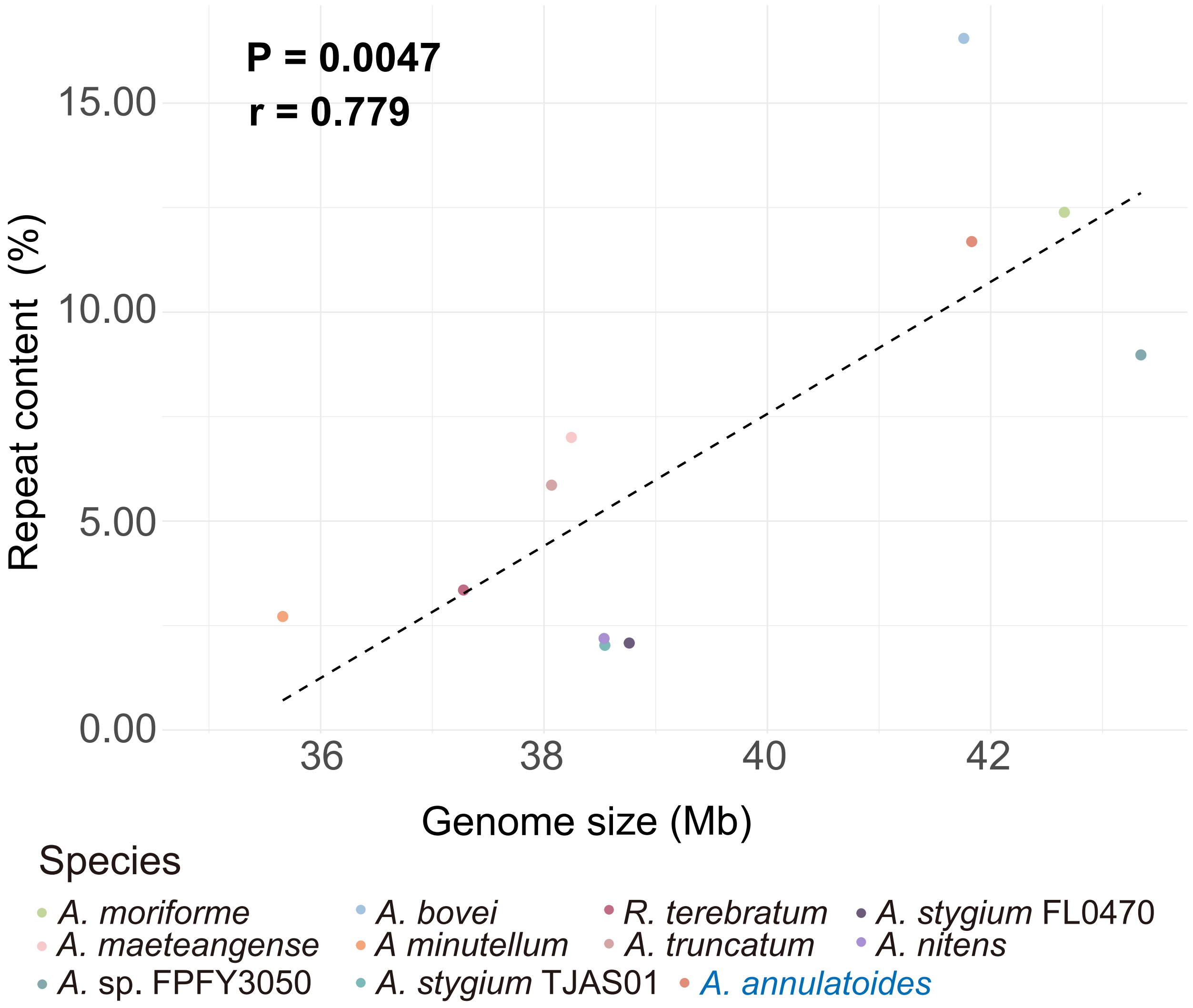


**Supplementary Figure 7. Genome size scales with repeat content across *Annulohypoxylon* species.** Scatterplot showing the relationship between total genome size (Mbp) and total repeat content (%) across 11 *Annulohypoxylon* species. Each point represents one species, coloured by taxon, with *A. annulatoides* highlighted for reference.

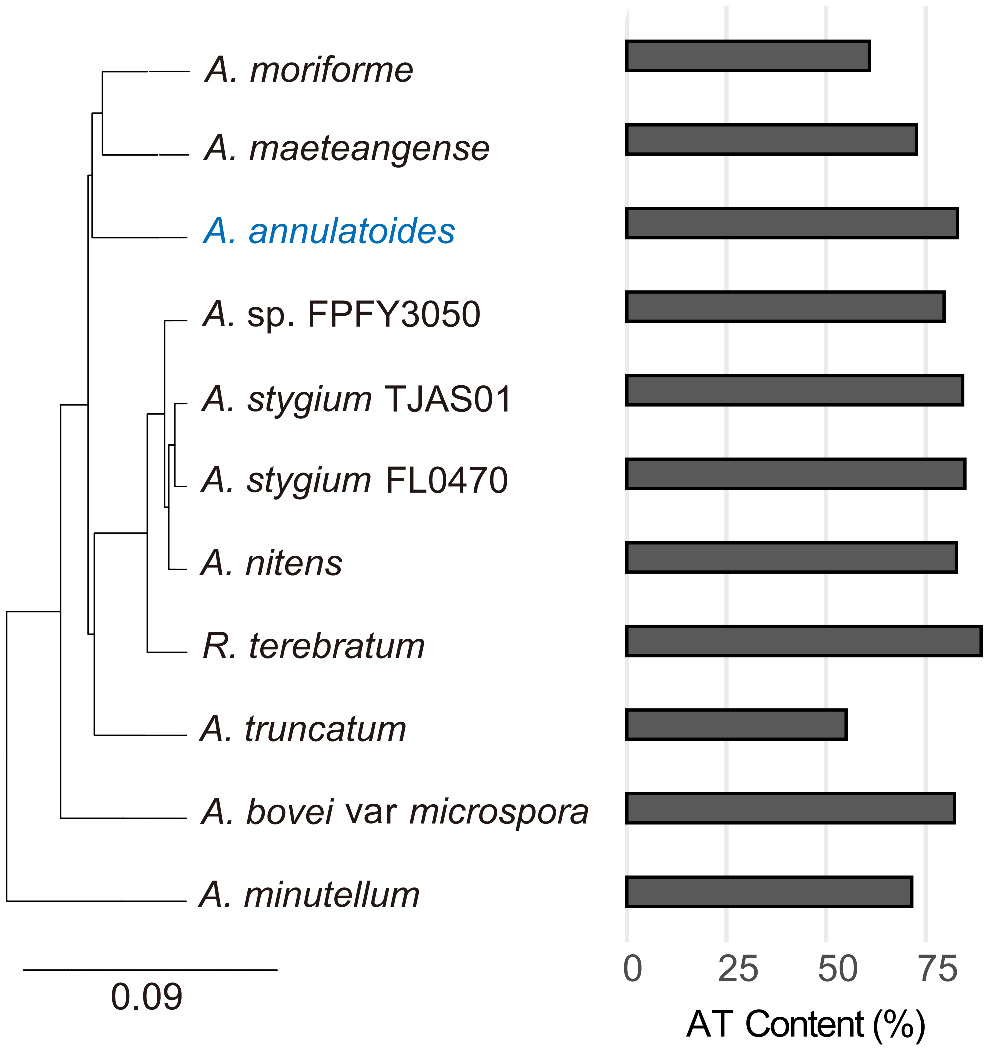


**Supplementary Figure 8. AT content of repeat sequences across the genomes of 11 *Annulohypoxylon* species.**

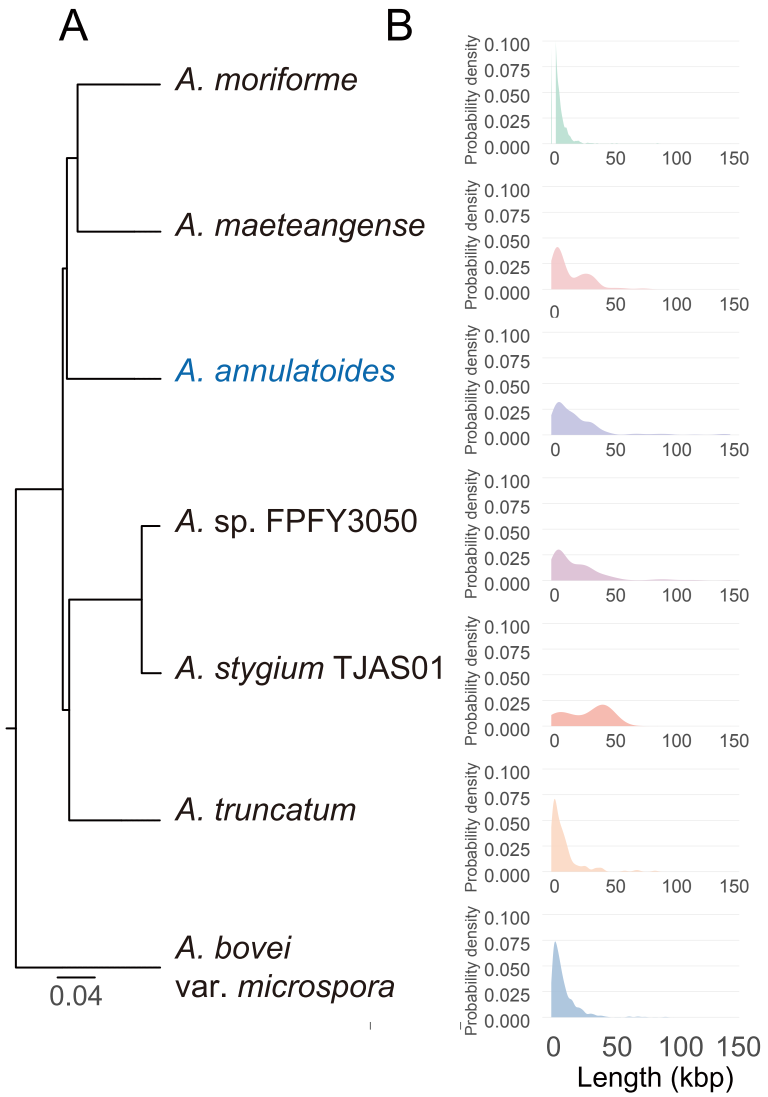


**Supplementary Figure 9. Length distributions of AT-rich isochores across *Annulohypoxylon* species.** (A) Phylogenetic relationships among the seven Hypoxylaceae species analysed, highlighting the placement of *A. annulatoides* relative to closely related taxa. (B) Kernel density estimates showing the length distributions of AT-rich isochores in *A. moriforme*, *A. maeteangense*, *A. annulatoides*, *A.* sp. FPYF3050, *A. stygium* TJAS01, *A. truncatum*, and *A. bovei* var. *microspora*.

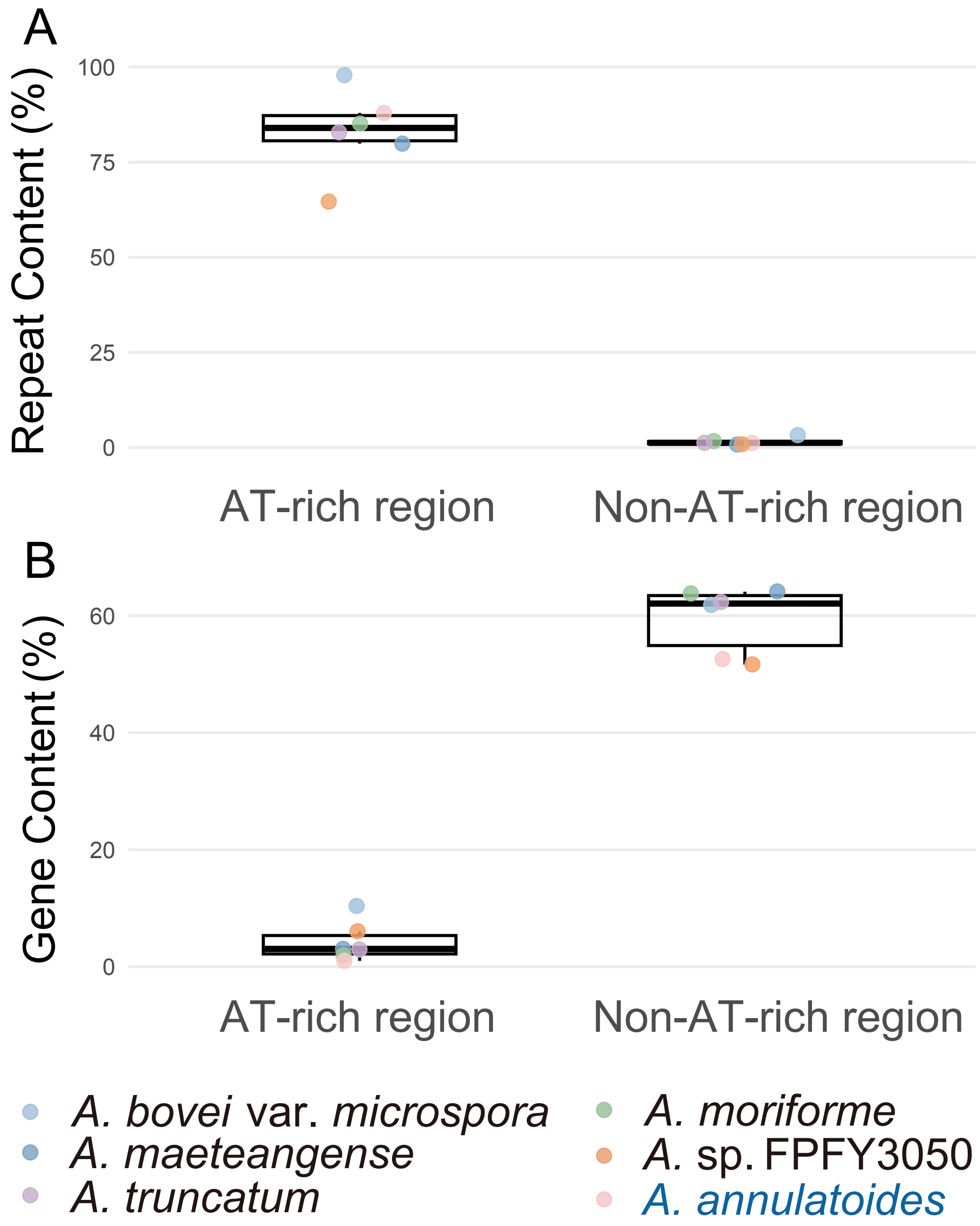


**Supplementary Figure 10.** Repeat coverage in regions with different AT content. (a) Comparison of coverage between genomic regions with AT content > 60 and (b) those with AT content < 60%.

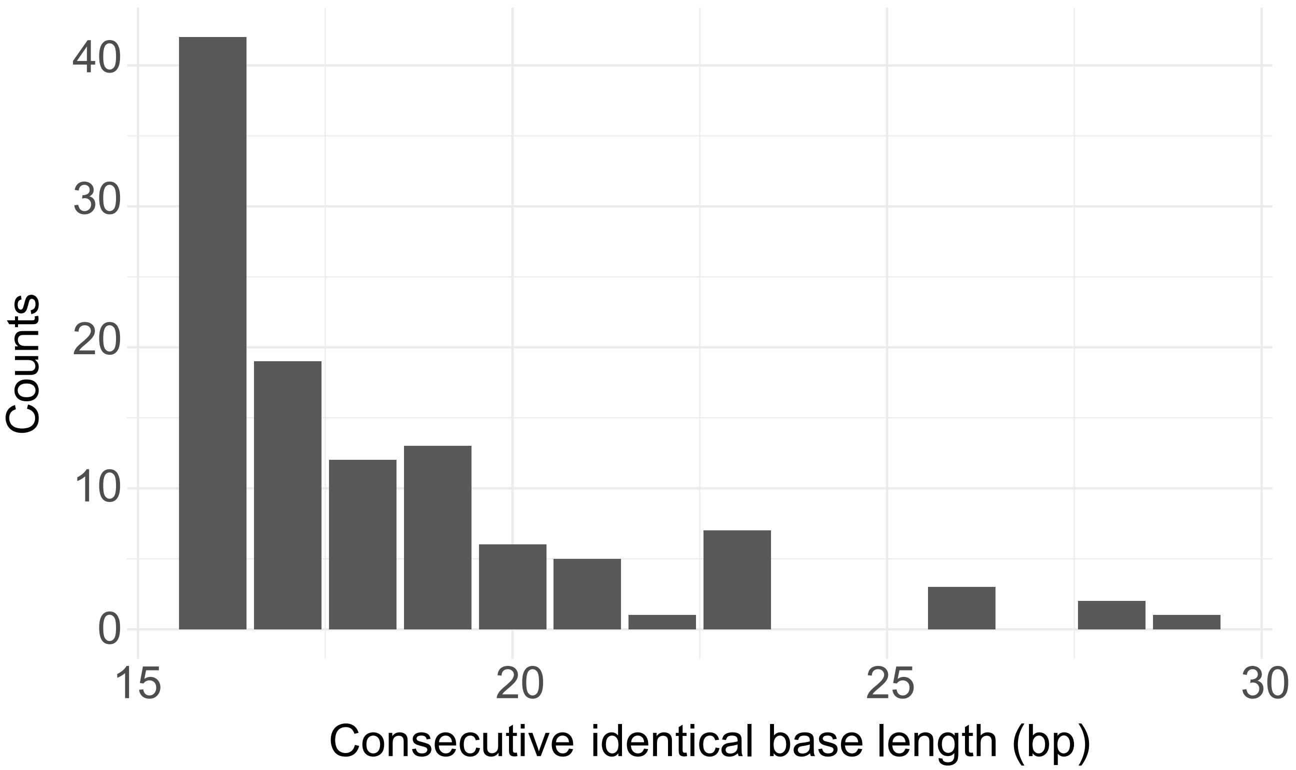


**Supplementary Figure 11. Distribution of homopolymers longer than 15 bp in AT-rich isochores.**

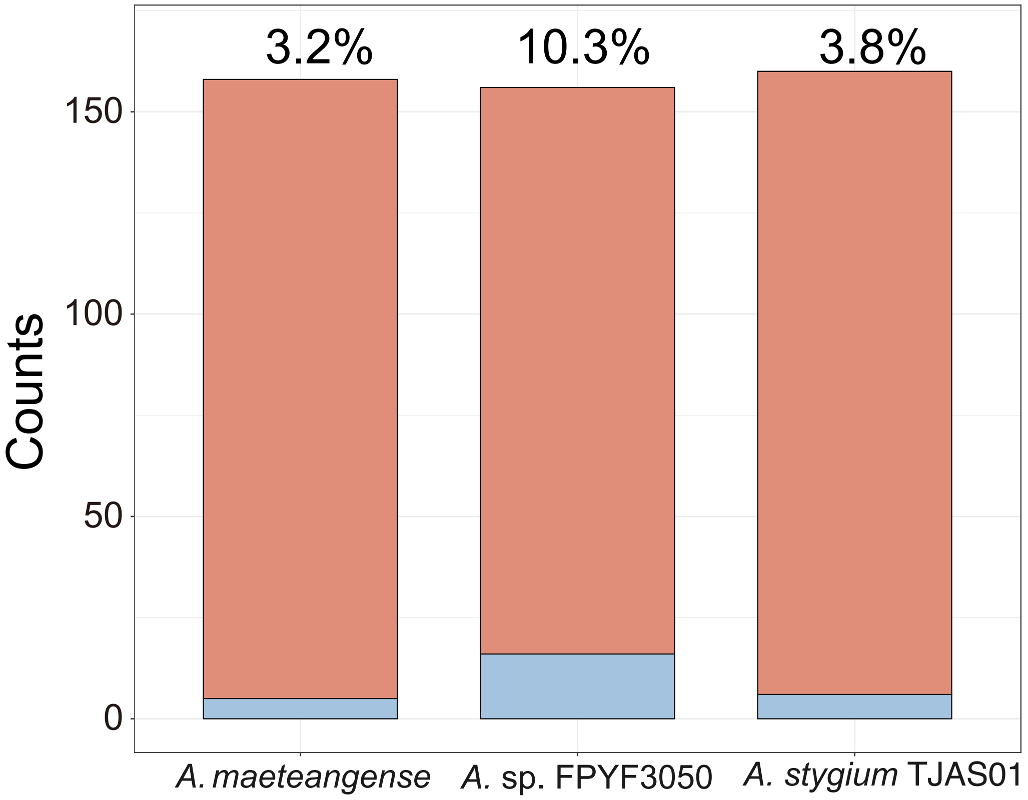


**Supplementary Figure 12. Limited positional conservation of AT-rich isochores across *Annulohypoxylon* species.** Stacked bar plots show intergenic regions in three *Annulohypoxylon* species (*A. maeteangense*, *A.* sp. FPYF3050, and *A. stygium* TJAS01) that either share an AT-rich isochore with *A. annulatoides* (blue) or lack an isochore at the corresponding position (salmon). Percentages above bars indicate the proportion of intergenic regions with shared isochores.

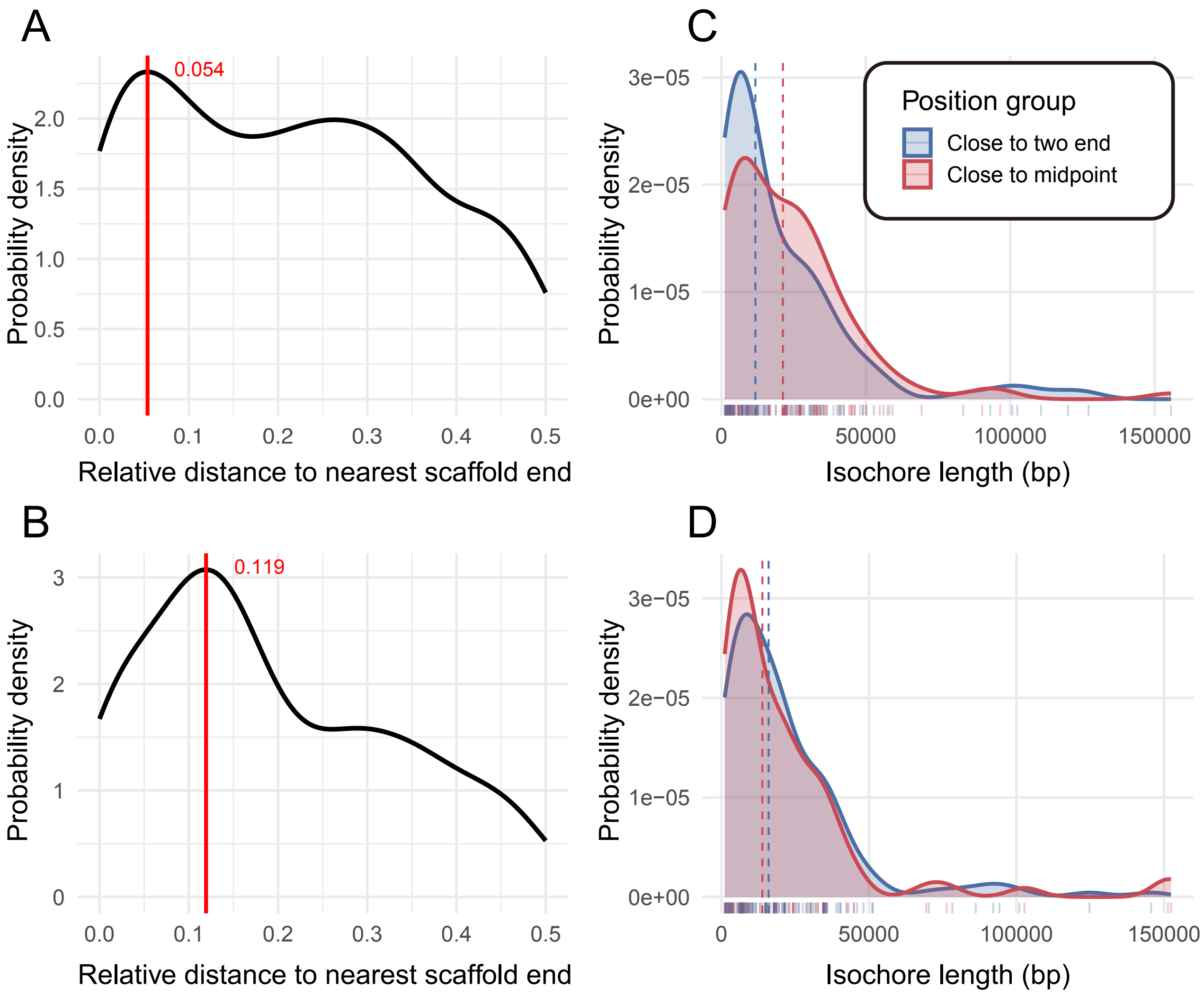


**Supplementary Figure 13. Length distributions of AT-rich isochores along chromosomes.** (A, B) Kernel density distributions showing the relative positions of AT-rich isochores in *A.* sp. FPYF3050 (A) and *A. annulatoides* (B), measured as the normalized distance to the nearest scaffold end (0 = scaffold end; 0.5 = scaffold midpoint). Vertical dashed lines indicate median positions, (C, D) Kernel density distributions of AT-rich isochore lengths in *A.* sp. FPYF3050 (C) and *A. annulatoides* (D), stratified by positional category. Isochores with relative distance ≤ 0.25 were classified as near-terminal, whereas those with distance > 0.25 were classified as central. Dashed lines mark median isochore lengths for each group.

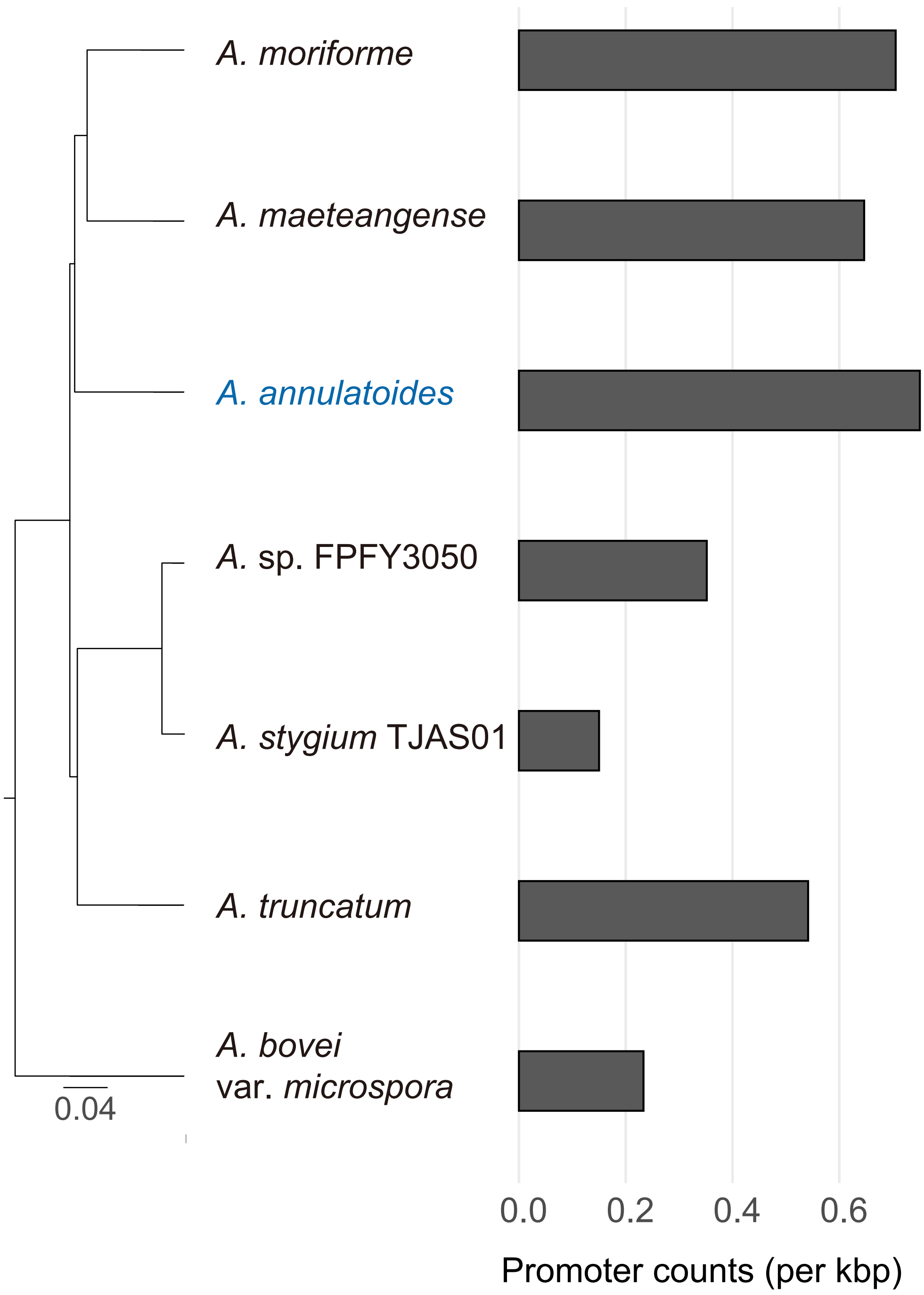


**Supplementary Figure 14. Enrichment of promoter-like motifs in AT-rich isochores across *Annulohypoxylon* species.** Bar plots show promoter density (predicted promoters per 1 kbp) within AT-rich isochore regions across *Annulohypoxylon* species, including *A. annulatoides*, *A. moriforme*, *A. maeteangense*, *A.* sp. FPYF3050, *A. truncatum*, and *A. bovei* var. *microspora*.

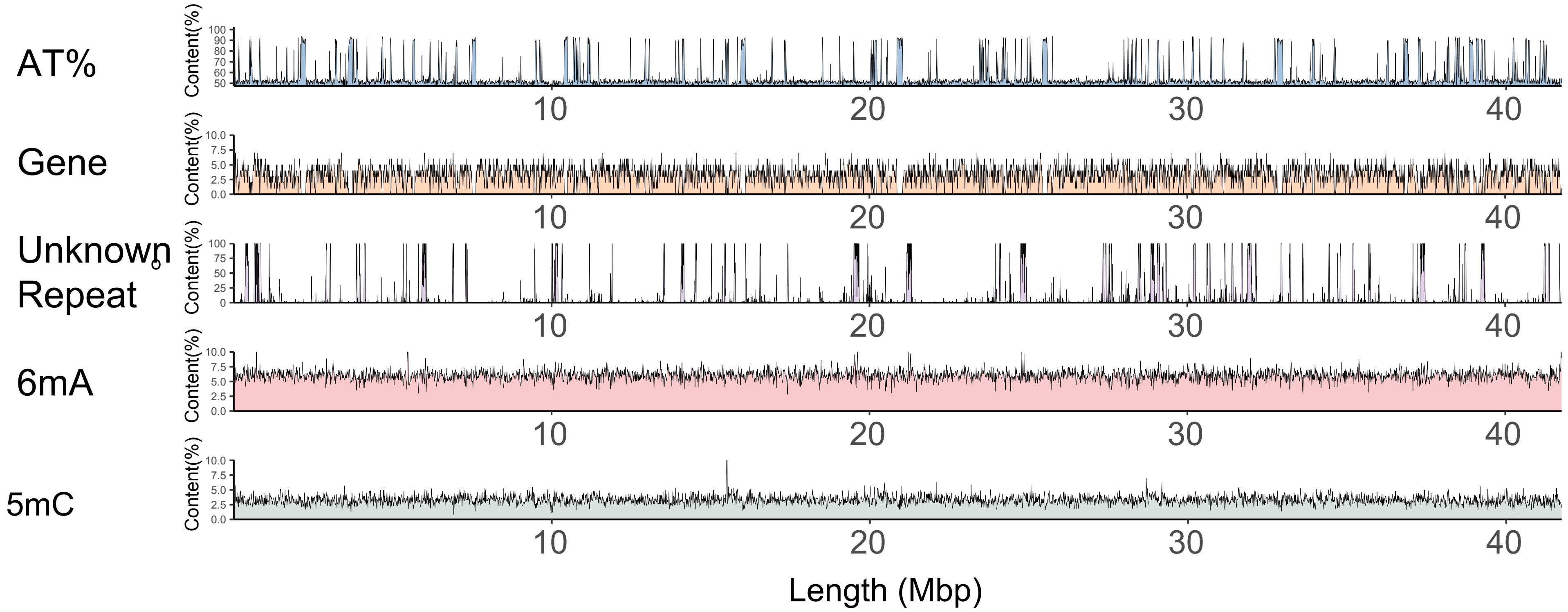


**Supplementary Figure 15. Genome-wide distribution of sequence composition and epigenetic features in *Annulohypoxylon annulatoides*.** Tracks show genome-wide profiles of gene density, repeat density, AT content, N6-methyldeoxyadenosine (6mA) methylation, and 5-methylcytosine (5mC) methylation across the *A. annulatoides* genome, calculated using a 10-kb sliding window.

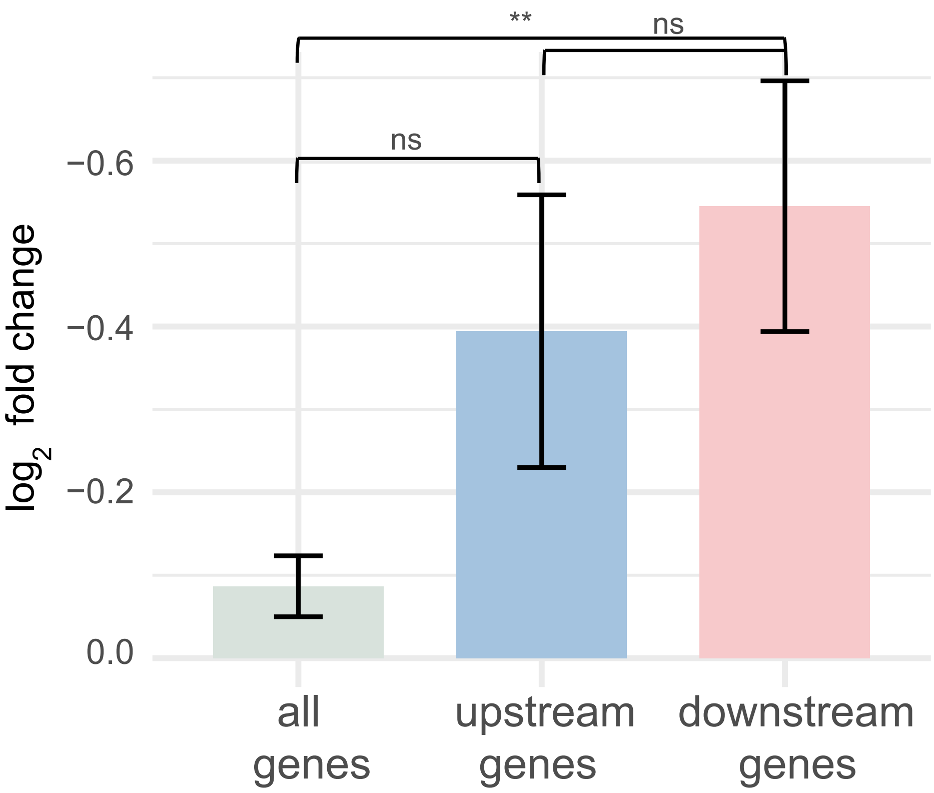


**Supplementary Figure 16. Expression of genes flanking AT-rich isochores under standard growth conditions.** Bar plots show the mean $\log_2$ relative expression of all genes (control), genes immediately upstream, and genes immediately downstream of AT-rich isochores in *Annulohypoxylon annulatoides*. Statistical significance was assessed using Student's t-test (ns, not significant; $^{**}P < 0.01$; $^{***}P < 0.001$).

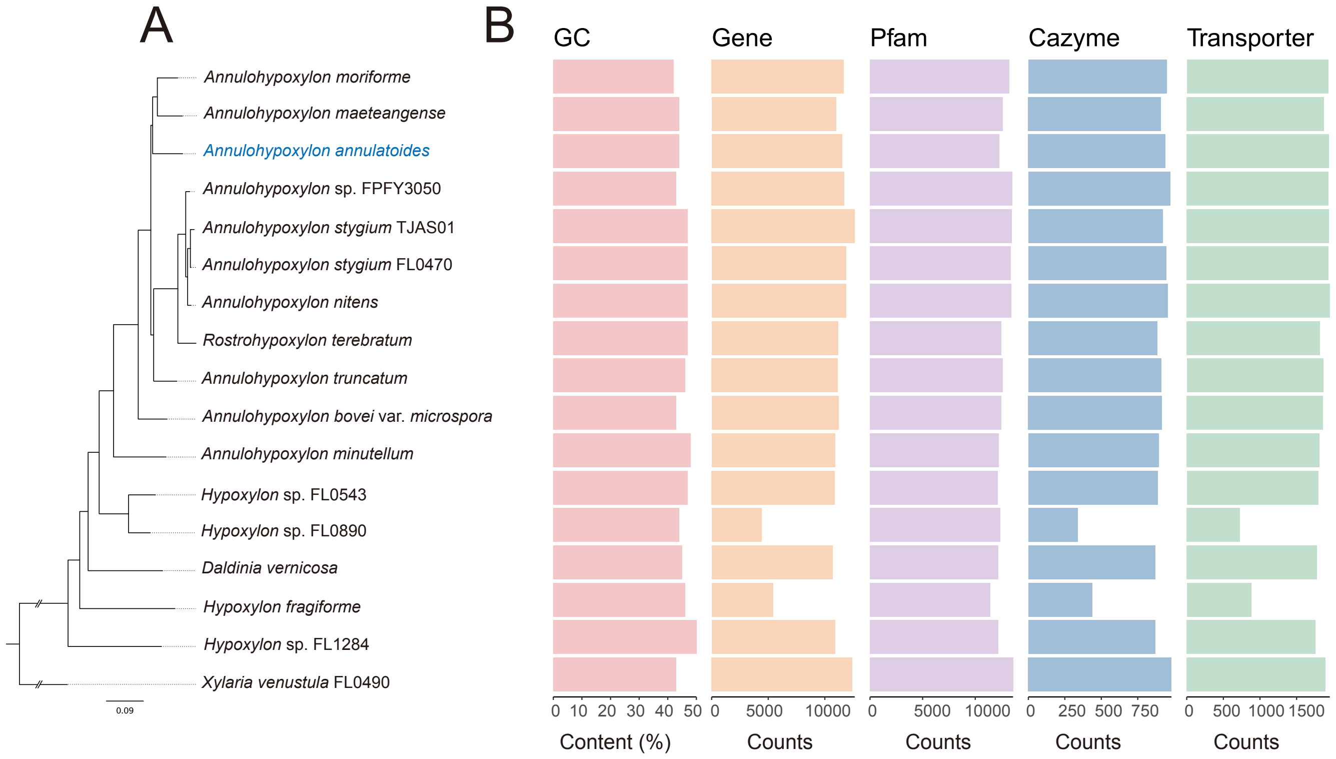


**Supplementary Figure 17. Conserved gene and functional repertoire across Hypoxylaceae genomes.** (A) Phylogenetic relationships among 16 Hypoxylaceae species with *Xylaria venustula* FL0490 shown as the outgroup; *Annulohypoxylon annulatoides* is highlighted for reference. (B) Comparative profiles of genome composition and functional gene content across species, including GC content (%), total gene counts, Pfam domain counts, carbohydrate-active enzymes (CAZymes), and predicted transporters.

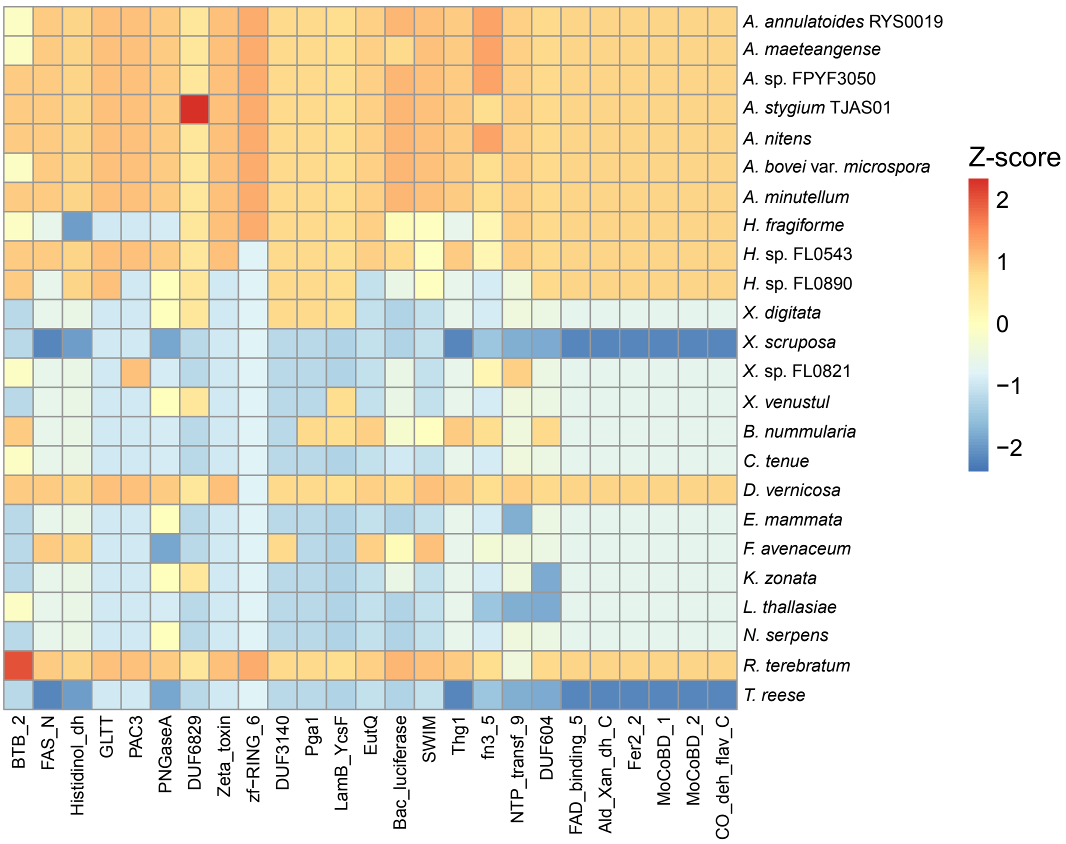


**Supplementary Figure 18. Protein families enriched in Hypoxylaceae relative to other ascomycete fungi.** Heatmap showing protein families that are significantly more abundant in Hypoxylaceae species compared with non-Hypoxylaceae ascomycete outgroups. Protein families were identified using a Wilcoxon rank-sum test with Benjamini–Hochberg correction (adjusted $P < 0.05$).
